# wQFM: Statistically Consistent Genome-scale Species Tree Estimation from Weighted Quartets

**DOI:** 10.1101/2020.11.30.403352

**Authors:** Mahim Mahbub, Zahin Wahab, Rezwana Reaz, M. Saifur Rahman, Md. Shamsuzzoha Bayzid

## Abstract

**Motivation:** Species tree estimation from genes sampled from throughout the whole genome is complicated due to the *gene tree-species tree discordance*. Incomplete lineage sorting (ILS) is one of the most frequent causes for this discordance, where alleles can coexist in populations for periods that may span several speciation events. Quartet-based summary methods for estimating species trees from a collection of gene trees are becoming popular due to their high accuracy and statistical guarantee under ILS. Generating quartets with appropriate weights, where weights correspond to the relative importance of quartets, and subsequently amalgamating the weighted quartets to infer a single coherent species tree allows for a statistically consistent way of estimating species trees. However, handling weighted quartets is challenging.

**Results:** We propose wQFM, a highly accurate method for species tree estimation from multi-locus data, by extending the quartet FM (QFM) algorithm to a weighted setting. wQFM was assessed on a collection of simulated and real biological datasets, including the avian phylogenomic dataset which is one of the largest phylogenomic datasets to date. We compared wQFM with wQMC, which is the best alternate method for weighted quartet amalgamation, and with ASTRAL, which is one of the most accurate and widely used coalescent-based species tree estimation methods. Our results suggest that wQFM matches or improves upon the accuracy of wQMC and ASTRAL.

**Availability:** wQFM is available in open source form at https://github.com/Mahim1997/wQFM-2020.

## 1 Introduction

With the rapid growth rate of newly-sequenced genomes, it is now common to estimate species trees from genes sampled throughout the whole genome. However, each individual gene has its own phylogeny (known as a gene tree), which may not agree with the species tree. Species tree estimation from multi-locus datasets is complicated in the presence of species tree-gene tree heterogeneity when gene trees differ, which can result from many biological processes, of which incomplete lineage sorting (ILS), modelled by the multispecies coalescent (MSC) [1], is probably the most common. ILS is also known as “deep coalescence”, which occurs with high probability whenever the time between speciation events is short relative to the population size [2]. ILS presents substantial challenges to species tree estimation [3,4]. For example, the standard approach, concatenation (which concatenates multiple sequence alignments of different genes into a single super-alignment and then estimates a tree from this alignment) can be statistically inconsistent [5] and can return incorrect trees with high confidence [6–9]. Moreover, under some conditions, the most probable gene tree topology may not agree with the species tree, which is known as the “anomaly zone” [3,10].

As a result of these studies, “summary methods”, which operate by computing gene trees from different loci and then combine the inferred gene trees into a species tree, are becoming increasingly popular [11], and many of these summary approaches are statistically consistent under the MSC model [12–23]. Using the most basic pieces of phylogenetic information (i.e., triplets in a rooted setting, and quartets in an unrooted setting) are key to the design of some of the statistically consistent methods [15,20,24]. ASTRAL, which is one of the most accurate and widely used coalescent-based methods, tries to infer a species tree so that the number of induced quartets in the gene trees that are consistent with the species tree is maximized. Another approach is to infer individual quartets, and then amalgamate these quartets into a single coherent species tree [20, 24–30].

Quartet amalgamation techniques have drawn substantial interest both from practical and theoretical perspectives as quartets can be inferred from raw data with high accuracy and these quartets can subsequently be amalgamated to obtain a highly accurate species tree [20, 25, 31–33]. The summary methods are usually sensitive to gene tree estimation error [11, 34, 35], so methods that can estimate species trees without needing to compute gene trees are of utmost importance. Therefore, quartet amalgamation techniques have attracted a lot of interest among the systematists. For example, SVDquartets [20] reliably estimates quartets from sequence data under the coalescent model using techniques from algebraic statistics, and then assemble these quartets to infer a species tree. However, a given set of quartets may not be *compatible* with a single species tree, and estimating a species tree by finding the largest compatible subset of a given quartet set is computationally hard [36]. Moreover, this approach implicitly gives the same importance to all quartets, and thus does not take the relative reliabilities of various quartets into account [27].

There is ample evidence that assigning weights to quartets (where weight of a quartet denotes the relative confidence of a particular quartet topology out of the three alternate topologies on a set of four taxa) can improve phylogenetic analyses [27,32,37]. This grow-ingawareness about the importance of weighted quartets has led to the development of methods like wQMC [30], which is an weighted extension of the quartet max-cut (QMC) algorithm [29]. In this paper, we present a new method for amalgamating weighted quartets by extending Quartet Fiduccia-Mattheyses (QFM) algorithm [24], which is being widely used in important phylogenetic studies [38–47], especially along with SVDquartets [20,48]. SVDquartets has been implemented in PAUP* [49], which uses QFM as a quartet agglomeration technique [50]. The proposed extension of QFM, weighted QFM (wQFM), is provably statistically consistent under the MSC. This is a divide-and-conquer based approach, and it introduces a novel scheme for defining the “partition score” to assess the quality of the candidate partitions during each divide step of the algorithm. We report, on an extensive evaluation study using widely used simulated dataset as well as biological dataset, the performance of wQFM. We reanalyzed a collection of biological dataset, including the avian phylogenomic dataset [51] comprising 14,446 genes across 48 genomes representing all avian orders. Our results suggest that wQFM achieves competitive or in some cases better tree accuracy than the main two alternatives, wQMC and ASTRAL. Notably, wQFM produced a relatively more reliable avian tree than ASTRAL.

## 2 Approach

wQFM uses a two-step technique in which we first use the input set of estimated gene trees to produce a set of weighted four-taxon trees (called “weighted quartet trees”, or “weighted quartets”), and then combine these weighted quartet trees into a tree on the full set of taxa using a heuristic aimed at finding a species tree of minimum distance to the set of weighted quartet trees (details below). Thus, wQFM is similar in overall structure to the population tree in BUCKy [18,52], and its proof of statistical consistency follows the same arguments as those provided for BUCKy-pop [52]. To understand wQFM, we begin by defining a very simple approach, Combining Dominant Quartet Trees 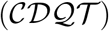, which is also statistically consistent. The proof that CDQT is statistically consistent explains why wQFM and BUCKy-pop are statistically consistent, and motivates their algorithmic designs.

### Combining Dominant Quartet Trees 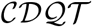

The basic idea is to take the input set of gene trees, compute a “dominant quartet tree” (see below) for every four species, and then combine the dominant quartet trees into a supertree on the full set of species using a preferred quartet amalgamation technique. If the quartet amalgamation technique correctly computes the supertree when the dominant quartet trees are “compatible” (see below), then CDQT is a statistically consistent method under the multi-species coalescent. Thus, CDQT depends on the quartet amalgamation technique, and so is a general technique, and not a particular technique.

The input to CDQT is a set 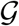 of unrooted gene trees, one for each gene, and each gene tree has the set S of species for its leafset. We denote by *ab|cd* the induced quartet on four species *a,b,c, d*, where the internal branch separates *a* and *b* from *c* and *d*. Note that on a set of four species *a, b, c, d*, there are three possible quartet trees (*ab|cd, ac|bd*, and *bc|ad*). The CDQT algorithm has the following steps:

1. For every set of four species *a, b, c, d*, we compute the set of induced four-leaf trees, one for each gene.
2. For every four leaves *a, b, c, d*, we determine which of the three possible unrooted trees on *a, b, c, d* occurs the most frequently; this is called the “dominant quartet tree on a, b, c, d”.
3. We construct a tree T from the set of dominant quartet trees using a preferred quartet amalgamation technique.

Because the method depends on the choice of quartet amalgamation technique, we now discuss this issue. We say that a set of quartet trees is “compatible” if there is a tree T such that every quartet tree is topologically identical to the subtree of T induced on its leaf set. Furthermore, when the set of quartet trees is compatible, then there is a unique tree *T*′ that induces all the quartet trees (called the “compatibility supertree”), and it can be computed in polynomial time using very simple techniques (e.g., the Naive Quartet Method, discussed in [53]). Thus, while there are many quartet amalgamation techniques, most of them are able to return the compatibility supertree when the input set contains a tree on every four leaves and is compatible. We call such quartet amalgamation techniques “proper”.

#### Theorem 1.

*If the quartet amalgamation technique is proper, then 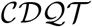 is statistically consistent under the multi-species coalescent model*.

#### Proof.

Let (*T*_0_, Θ) be an unrooted model species tree in the multi-species coalescent model (and so *T*_0_ is a binary species tree and Θ are the branch lengths in *T*_0_ in coalescent units). Let *Q* be a set of *k* gene trees sampled under the multispecies coalescent model on (*T*_0_, Θ). By [10], for every four species *a, b, c, d* (leaves in *T*_0_), the most probable unrooted gene tree on *a, b, c, d* is topologically identical to the unrooted tree induced by *T*_0_ on *a, b, c, d*. Therefore, when *k* (the number of gene trees) is large enough, with high probability the dominant quartet tree on *a, b, c, d* will be equal to the species tree on *a, b, c, d*. Hence, for large enough *k*, with high probability the set of dominant quartet trees will be compatible, and will uniquely identify the unrooted species tree; when this holds, 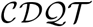 will reconstruct the species tree. In other words, for every *ϵ* > 0, there is some value *K* so that for *k > K* and given *k* true gene trees sampled from the probability distribution on true gene trees defined by (*T*_0_, Θ), with probability at least 1 − *ϵ*, the dominant quartet trees will be equal to the induced four-leaf species trees, and any proper quartet tree amalgamation technique will correctly reconstruct the species tree. Hence, 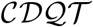 is statistically consistent under the multi-species coalescent model.

The proof that 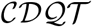 is statistically consistent under the multispecies coalescent model provides a guarantee under idealized conditions – where all gene trees are correct and there are a sufficiently large number of them. However, in practice estimated gene trees have error and there may not be a sufficiently large number of gene trees. Therefore, for good performance (and not just theoretical guarantees), species tree estimation methods need to work well with estimated gene trees – for which the dominant gene trees may not be identical to the most probable gene trees, and hence may not be compatible with each other. Therefore, heuristics for combining quartet trees, that can construct supertrees even when the quartet trees are incompatible, are valuable techniques for species tree estimation in the presence of ILS.

### Related methods

BUCKy-pop [18] (the population tree output by BUCKy) is one of the statistically consistent methods for species tree estimation under the multi-species coalescent model that uses a quartet-based approach. The input to BUCKy is a set of unrooted gene tree distributions, with one distribution per gene. (BUCKy was originally intended for use with posterior distributions computed using Bayesian MCMC methods, but has also been used with distributions computed using maximum likelihood bootstrapping; both approaches give similar results [54]). In the first step, BUCKy-pop uses the gene tree distributions to estimate a quartet tree for every four species, and performs this estimation using Bayesian techniques. In the second step, it combines these estimated quartet trees using a quartet tree amalgamation technique [28]. Because the quartet tree amalgamation technique will reconstruct the compatibility supertree if it exists, BUCKy-pop is statistically consistent under the multi-species coalescent model. Another popular method is SVDquartets [20, 48], where the loci in a multi-locus dataset are concatenated into a single long alignment, and then, for each set of four species, a quartet tree for that set is computed using algebraic statistics and singular value decomposition (SVD). Finally, a species tree is estimated (using QFM or QMC) by amalgamating the quartet trees so that it agrees with as many of these quartet trees as possible. ASTRAL is another quartet based method, which explores the tree space under a dynamic programming based algorithm which uses a weighted quartet score of a candidate species tree defined to be the number of quartets from the set of input gene trees that agree with the candidate species tree.

### wQFM

We present a new summary method, wQFM, which estimates species trees by combining gene trees. However, unlike SVDquartets, BUCKy-pop and 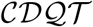, which consider a single quartet for each set of four taxa and compute species trees by combining these “dominant” quartet trees, wQFM computes weights for every possible four-leaf tree (and so for each of the three possible unrooted trees for every four leaves), and then combines this set of weighted quartet trees into a tree on the full set of species. However, wQFM can also be used to amalgamate a set of dominant quartets. In this study, we consider the frequency of a quartet, in the input gene trees, as its weight and so high weight suggests higher confidence in the quartet gene tree. However, weights can be inferred in other ways as well. For example, pairwise distances between taxa can be considered while computing the weights on quartets [30]. The weights can be computed from input set of gene trees with or without generating bootstrapping gene tree samples. wQFM uses a heuristic to combine the weighted quartet trees into a supertree, attempting to solve a version of the NP-hard “Maximum Quartet Compatibility” problem [55], where we set weights on the quartet trees.

### 2.1 Algorithmic pipeline of wQFM

wQFM has two steps; in the first step, we generate a set of weighted quartet trees from the input, and in the second step we estimate the species tree from the set of weighted quartet trees. We now describe how we perform each step.

#### Step 1: Generate weighted quartets

Given a set 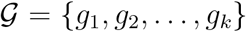 of *k* gene trees on taxon set 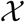, we compute weights for every possible quartet tree *ab|cd* of four leaves, where *ab|cd* denotes the unrooted quartet tree with leaf set 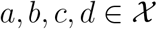 in which the pair *a, b* is separated from the pair *c, d* by an edge. Thus, we compute a weight *w*(*q*) for every possible (unrooted) quartet tree *q*. Note that on every set of four species, there are three possible unrooted quartet trees (simply called “quartets”). Also note that every gene tree on the set 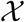 of taxa induces a single quartet tree on *a, b, c, d*. We define the support for quartet tree *ab|cd* to be the number of the trees in 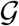 that induce *ab|cd* on set *a, b, c, d*.

#### Step 2: Construct supertree

Our technique, which we call wQFM, to combine the quartet trees into a tree on the full set of taxa is the weighted version of the QFM technique, developed in Reaz *et al*. [24]. We briefly describe the algorithmic pipeline, and refer the readers to Reaz *et al*. [24] for more details.

##### Terminology

For an unrooted tree *T* on taxon set 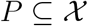, we let *L*(*T*) denote the leaf set of *T*. Every edge in *T* defines a bipartition of its leaf set (defined by deleting the edge but not its endpoints from *T*), which is denoted by *π_e_*. However, we can also refer to an arbitrary bipartition on set *P*, whether or not it is present in a given tree *T*; thus, we let (*X, Y*) be a bipartition with *X* on one side and *Y* on the other (note that the order of *X* and *Y* does not matter).

Under the assumption that all gene trees in the input are fully resolved, then given a bipartition (*X, Y*) on set 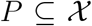, we partition the quartet trees defined by the input trees as follows:

- quartet trees that are *satisfied* by (*X, Y*): those quartet trees *ab|cd* where {*a, b*} ⊆ *X* and {*c, d*} ⊆ *Y*, or {*a, b*} ⊆ *Y* and {*c, d*} ⊆ *X* (i.e., the bipartition (*X, Y*) separates the two sibling leaf pairs with the quartet tree from each other),
- quartet trees that are *violated* by (*X, Y*): those quartet trees *q* whose taxa are fully contained in *X*∪*Y*, and where *X* and *Y* each contains exactly two of the four taxa in *q* but *q* is not satisfied by (*X, Y*), and
- quartet trees that are *deferred* by (*X, Y*): those quartet trees *q* so that ≥3 of its four taxa reside in *X* or in *Y*.

In fact, we can partition all possible quartet trees using any given bipartition, whether or not they appear in any input gene tree. We will refer to a pair (*X, Y*) with 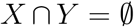 and 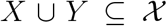 as either a full bipartition (or simply a bipartition). A non-trivial bipartition is one that has at least two taxa on each side.

### 2.2 Divide-and-conquer approach

Let *Q* be a set of weighted quartet trees over a taxon set *P ⊆ S*. The divide-and-conquer approach takes the pair (*Q, P*) as input. The basic divide-and-conquer algorithm operates in a top-down manner: a good nontrivial bipartition is produced, rooted trees are calculated on the two parts of the bipartition, and then combined together into a rooted tree on the full dataset by making them both subtrees of a common root. Then the tree is unrooted. The key to the algorithm is therefore finding the bipartition, and showing how to recurse on the subproblems so as to produce rooted trees.

This is the same basic top-down technique as used in Quartets MaxCut (QMC) [29], so the only difference in the two methods is how the good non-trivial bipartition is produced. The differences in algorithm design define how the tree space is being searched and are important to the accuracy of the resultant tree.

We now briefly describe the technique used to find a good bipartition. We score a bipartition with respect to the set *Q* of quartet trees based on the total weight of all satisfied quartets and the total weight of all violated quartets. However, the partition score can be defined in other ways and the number of deferred quartets can be considered as well [24]. We have proposed a new approach, where a *weighted* difference of the total weight of all satisfied quartets and the total weight of all violated quartets is used as a partition score. These weights are computed based on the distribution of the weights of the quartets (see Section “Partition score computation” for details). The technique to find the bipartition uses a heuristic iterative strategy, in which each iteration begins with the bipartition from the previous iteration, and tries to improve it. If the search strategy within this iteration finds a better bipartition, then a new iteration begins with the new bipartition. Thus, the strategy continues until it reaches a local optimum. The search within each iteration, however, allows for bipartitions with poorer scores to be computed, and hence the overall strategy is not purely hill-climbing. The running time of each iteration is polynomial, but the number of iterations depends on the search.

Given the final bipartition (*A, B*) on *P*, we use it to define two inputs to wQFM. By running wQFM recursively, we construct two rooted trees, one on *A* and one on *B*. We then create a rooted tree on *A ∪ B* (the full set of taxa), and then ignore the rooting to obtain an unrooted tree on 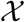. Thus, the rest of the algorithm depends on how we define these two inputs, and how we use wQFM to obtain rooted trees.

Letting (*A|B*) denote the bipartition that is produced in the divide step, we divide *Q* into three sets, as follows. The first set contains all quartet trees that are either satisfied or violated by (*A|B*). The other two sets are *Q_A_* and *Q_B_*, where *Q_A_* = {*q* ∈ *Q*: |*q ∩ A*| ≥ 3}, and *Q_B_* is defined similarly. Note that all quartet trees in *Q_A_* and *Q_B_* are deferred by (*A|B*).

For each quartet tree *q ∈ Q_A_* with |*q ∩ A*| = 3, we label the taxon that is not in *A* by a new dummy taxon *b*\*. We similarly relabel one leaf in the relevant quartet trees in *Q*_B_ with a new dummy taxon *a*\*. This produces sets 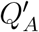 and 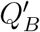, which are on sets 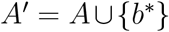 and 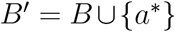, respectively. We then recurse on each pair 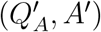 and 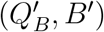, producing trees that we combine by identifying leaves *a*\* and *b*\*, and suppressing nodes of degree two. The base case is obtained when the taxon set has three or fewer leaves, in which case we return a star. We do not pursue this further here, but see Reaz *et al*. [24].

#### Theorem 2.

*wQFM is statistically consistent under the multispecies coalescent model*.

#### Proof.

Statistical consistency for a summary method follows if the method will return the true species tree given a sufficiently large number of true gene trees sampled from the distribution defined by the model species tree. So suppose we are given a large number of true gene trees so that the most probable gene tree is also the dominant gene tree. Therefore, the weights on quartet trees will be equal to the proportion of the gene trees that induce that particular quartet tree. Note that the weight of the dominant quartet tree will be greater than all other quartet trees. Because that the dominant quartet tree (one with the highest frequency) is the most probable gene tree, and hence also the true species tree for its leaf set (since there are no anomalous 4-leaf unrooted gene trees), the best score is obtained by the true species tree.

### 2.3 Partition score computation

For unweighted quartet amalgamation techniques, where, for each set of four taxa, we are given with one of the three alternate quartet topologies (as in SVDquartets), defining the partition score as the difference between the numbers of satisfied and violated quartets is a suitable criteria [24]. However, under the weighted setting – where all possible quartets are considered with relative weights – if one quartet topology on a set of four taxa is satisfied, the other two are guaranteed to be violated. Therefore, unless the weights of the satisfied quartets is substantially higher than the other two, the partition score will be negative. Hence, a partition score that assigns a relatively higher importance to the satisfied quartets is more reliable for weighted quartets. Let *w_s_* and *w_v_* be the total weights of the satisfied and violated quartets, respectively. In general, we can use *αw_s_ − βw_v_* as a partition score such that *αw_s_ − βw_v_* ≥ 0, where 0 < *α, β* ≤ 1. When only the dominant quartets are given as input (i.e., only one quartet for each set of four taxa), *α = β* = 1. Otherwise, we find appropriate values for *α* and *β* using an empirically selected heuristic, which takes the distribution of the weights of the quartets into account.

Let 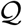 be the set of all possible sets of four taxa (for a set of *n* taxa, 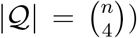. For the first divide step (where all the input quartets are considered), if the weights of the quartets are “closer” to each other on a reasonably small fraction of the four taxa sets in 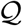, we set *α = β* = 1. Otherwise, we assign a relatively higher importance to the satisfied quartets by setting *α* = 1 and choosing an appropriate value for *β* using a heuristic approach. For a set 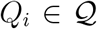, let *q*_1_, *q*_2_ and *q*_3_ be the three alternate quartet topologies with weights *w*_1_, *w*_2_, *w*_3_, respectively. Without loss of generality, we assume that *w*_1_ ≥ *w*_2_ ≥ *w*_3_. Setting *αw*_1_ − *β*(*w*_2_ + *w*_3_) = 0, and *α* to 1, the upper limit of *β* becomes equal to a ratio *r*, where 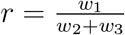. The minimum value of *r* is 0.5 (when three alternate quartets have equal weights, i.e., *w*_1_ = *w*_2_ = *w*_3_), and the value of *r* increases as the differences in the weights are increased. We compute *r* for each 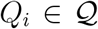. Next, we compute the fraction *f* of 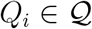, for which *r* lies within the range [0.5, λ), where λ is a empirically selected threshold. For the very first divide step, if *f* is relatively small (i.e., *f < δ* where *δ* is a user defined hyper parameter), we set *α* = *β* = 1, and use these values of *α* and *β* for all subsequent divide steps (i.e., we do not compute *β* heuristically in subsequent steps). Otherwise, we find a suitable value for *β* heuristically in every divide step. We have empirically selected *λ* = 0.9 and *δ* = 0.1. Note that different sets of quartets will be considered in different divide steps in the divide-and-conquer algorithm, and subsequently *β* will be changed accordingly.

We now describe the heuristic which we use to choose *β*. When *f ≥ δ*, we consider a series of bins of size 0.01 within the range [0.5, λ), i.e., *b*_1_ = [0.50, 0.51), *b*_2_ = [0.51, 0.52), …, *b*_(λ−0.5)∗10_ = [λ − 0.01, λ). If *f < δ* (not in the very first divide step, but in any of the subsequent ones), we consider a series of bins of size 0.01 within the range [λ, 1]. Next, for each bin *b_i_*, we compute *c_i_*, which denotes the number of 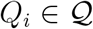 for which *r* lies in *b_i_*. We replace all the *r* values which are larger than 1 by 1 since very large *r* can inflate the overall computation. Finally, letting *m_b_i__* denote the mid-point of a bin *b_i_*, we compute *β* according to Eqn. 1. The pseudo-code of this heuristic algorithm is provided in supplementary materials.

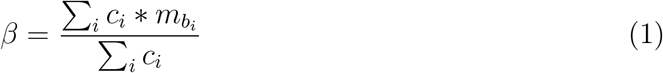

The running time of wQFM (and similar divide-and-conquer methods) depends on the rate of convergence to a good bipartition in every divide step. This, in turn, depends on the partition score which is used to evaluate a bipartition. See supplementary materials for additional results showing the impact of different partition scores on tree accuracy.

## 3 Experimental Studies

### 3.1 Datasets

#### Simulated dataset

We studied previously used simulated and biological datasets to evaluate the performance of wQFM. We used two biologically-based simulated datasets (the avian and mammalian simulated datasets) studied in [56]. We also analyzed three other simulated datasets (11-taxon, 15-taxon and 101-taxon) from [56–58]. These dataset range from moderately low to extremely high levels of ILS, and range in terms of gene tree estimation errors and numbers of genes.

This mammalian dataset was simulated by taking the species tree estimated by MP-EST on the biological dataset studied in Song *et al*. [59]. This species tree had branch lengths in coalescent units, which we used to produce a set of gene trees under the coalescent model. Thus, the model tree has an ILS level based on a coalescent analysis of the biological mammalian dataset, and other properties of the simulation that are set to reflect the biological sequences they studied. We explored the impact of varying numbers of genes (25 ~ 800), varying amounts of gene tree estimation error (i.e., the amount of phylogenetic signal by varying the sequence length for the markers: 250bp ~ 1500bp). In both cases, the levels of ILS were varied (shorter branches increases ILS) by multiplying or dividing all internal branch lengths in the model species tree by two. Thus, we have three model conditions that are referred to as 1X (moderate ILS), 0.5X (high ILS) and 2X (low ILS). The 48-taxon avian simulated dataset is based on the species tree estimated using MP-EST on the avian dataset of [51], and was simulated by following a similar procedure as the mammalian dataset. Similar to the mammalian dataset, it has three different ILS levels (1X, 0.5x and 2X), albeit the ILS levels are higher than the mammalian dataset (i.e., more discordance between the true gene trees and the species tree).

We analyzed the high-ILS 11-taxon datasets from [57] (as the model condition with lower amount of ILS is very easy to analyze [11]) which varies in the number of genes and amount of gene tree estimation error. 15-taxon datasets contain a high level of ILS and vary in sequence lengths and numbers of genes. Thus, the simulated datasets provide a wide range of challenging and practical model conditions in which we explore the performance of wQFM.

#### Biological dataset

We analyzed a collection of biological datasets: the 37-taxon mammalian dataset from Song *et al*. [59], the avian phylogenomic dataset containing 48 species and 14,446 loci (including exons, introns and UCEs), the amniota dataset from Chiari *et al*. [60], and the angiosperm dataset from Xi *et al*. [61].

### 3.2 Methods

We compared wQFM with the best existing weighted quartet amalgamation method wQMC as well as with ASTRAL-III [62] (version 5.7.3), which is considered as one of the most accurate and widely used coalescent-based species tree estimation methods. ASTRAL has been shown to outperform other coalescent based methods [15, 50, 63], including MP-EST, Bucky, NJst and SVDquartets. We ran wQFM and wQMC using the embedded quartets in the gene trees with weights reflecting the frequencies of the quartets. ASTRAL augments the set of bipartitions present in the input gene trees (by adding extra bipartitions) in order to search a larger tree space [58], and thus it extends the set of quartets in the input set of gene trees. This expansion resulted in an improved accuracy of ASTRAL-II over ASTRAL-I [58]. Therefore, the search space explored by ASTRAL could be larger than those considered by wQFM and wQMC.

### 3.3 Measurements

We compared the estimated trees (on simulated datasets) with the model species tree using normalized Robinson-Foulds (RF) distance [64] to measure the tree error. The RF distance between two trees is the sum of the bipartitions (splits) induced by one tree but not by the other, and vice versa. All the trees estimated in this study are binary and so False Positive (FP), and False Negative (FN) and RF rates are identical. For the biological dataset, we compared the estimated species trees to the scientific literature. We analyzed multiple replicates of data for various model conditions and performed Wilcoxon signed-rank test (with *α* = 0.05) to measure the statistical significance of the differences between two methods.

## 4 Results and Discussion

### 4.1 Results on 37-taxon dataset

The average RF rates of wQFM, wQMC and ASTRAL on various model conditions in 37-taxon dataset are shown in Fig. 1. Overall, ASTRAL, wQFM and wQMC had competitive accuracy, but wQFM achieved statistically significant improvement over Astral and wQMC on a few model conditions. The impact of changes in the ILS level with 500bp sequences and 200 genes is shown in Fig. 1(a). As expected, species tree error rates increased as ILS levels increased. ASTRAL and wQMC had similar accuracy, and wQFM was slightly better than these two existing methods (the differences are statistically significant (*p* ≤ 0.05) on 1X model condition). Figure 1(b) shows the impact of varying amounts of gene tree estimation error (controlled by sequence lengths). All methods showed improved accuracy as the sequence length was increased, and best results were obtained on true gene trees. wQFM consistently produced slightly better species trees than ASTRAL and wQMC, and its improvement over ASTRAL and wQMC is statistically significant on 250bp and 500bp model conditions. Fig. 1(c) shows that all these methods improved as we increased the number of genes, which is expected for statistically consistent methods. wQFM matched the accuracy of ASTRAL and wQMC with slight advantage on model conditions with relatively larger numbers of genes.

**Figure 1:**
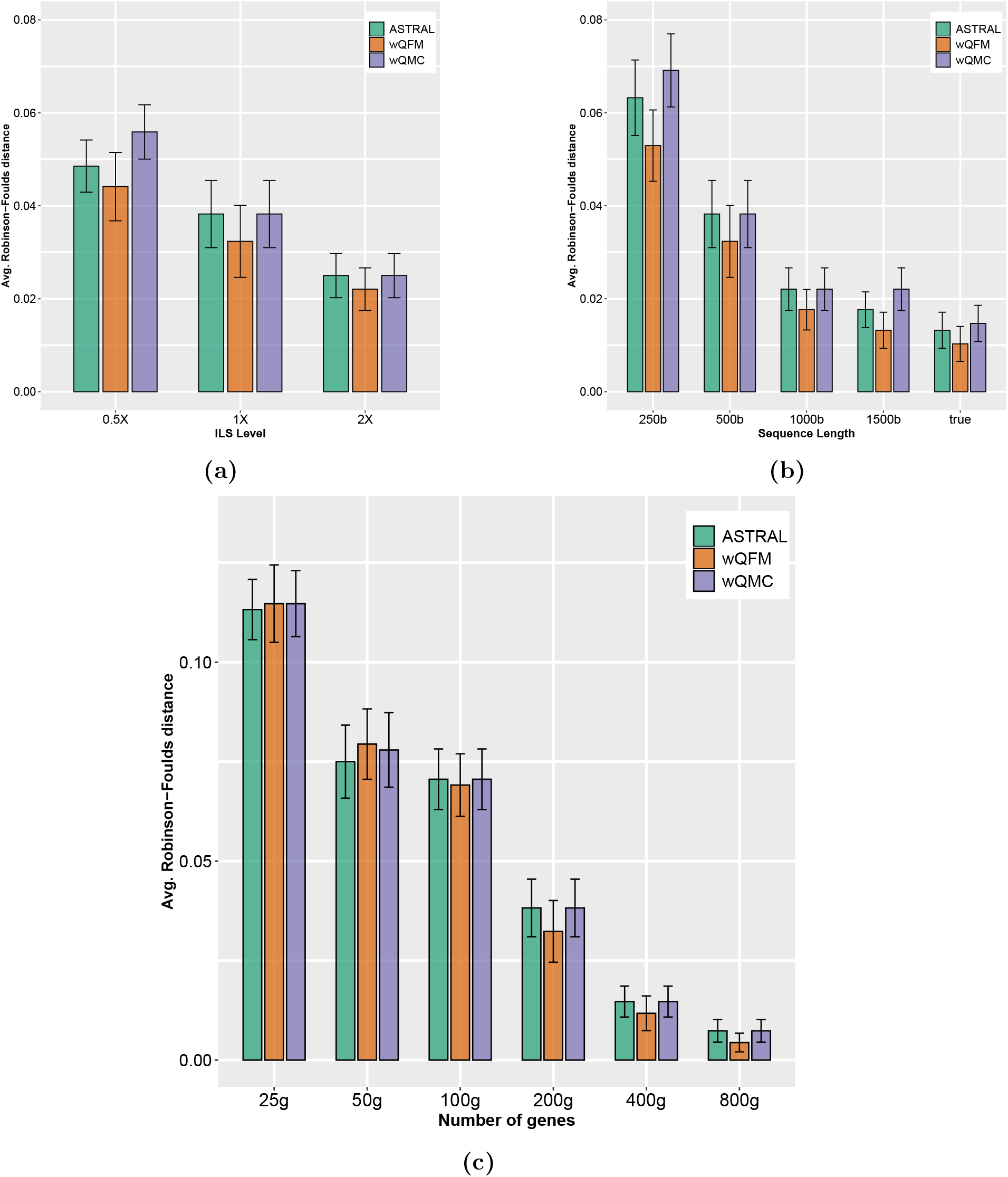
Comparison of ASTRAL, wQFM and wQMC on 37-taxon simulated mammalian dataset. We show the average RF rates with standard error bars over 20 replicates. (a) The level of ILS was varied from 0.5X (highest) to 2X (lowest) amount, keeping the sequence length fixed at 500bp and the number of genes at 200; (b) The sequence length was varied from 250bp to 1500bp, keeping the number of genes fixed at 200, and ILS at 1X (moderate ILS); (c) The number of genes was varied from 25g to 800g, with 500bp sequence length and moderate (1X) ILS.

### 4.2 Results on 11-taxon dataset

The performance of various methods on 11-taxon high-ILS dataset with varying numbers of estimated and true gene trees is shown in Fig. 2. On this dataset, similar to the 37-taxon dataset, ASTRAL and wQMC had very close performance, and wQFM was better than them in some cases (although the improvement was not always statistically significant). As was expected, the accuracy of these methods improved with the increase in the number of genes and they returned highly accurate species trees when true gene trees were used (even with only 25 genes). wQFM outperformed both ASTRAL and wQMC on small numbers of true gene trees (e.g., 5 and 15 gene model conditions), and returned the true species tree for all of the 20 replicates of data with 25 true gene trees.

**Figure 2:**
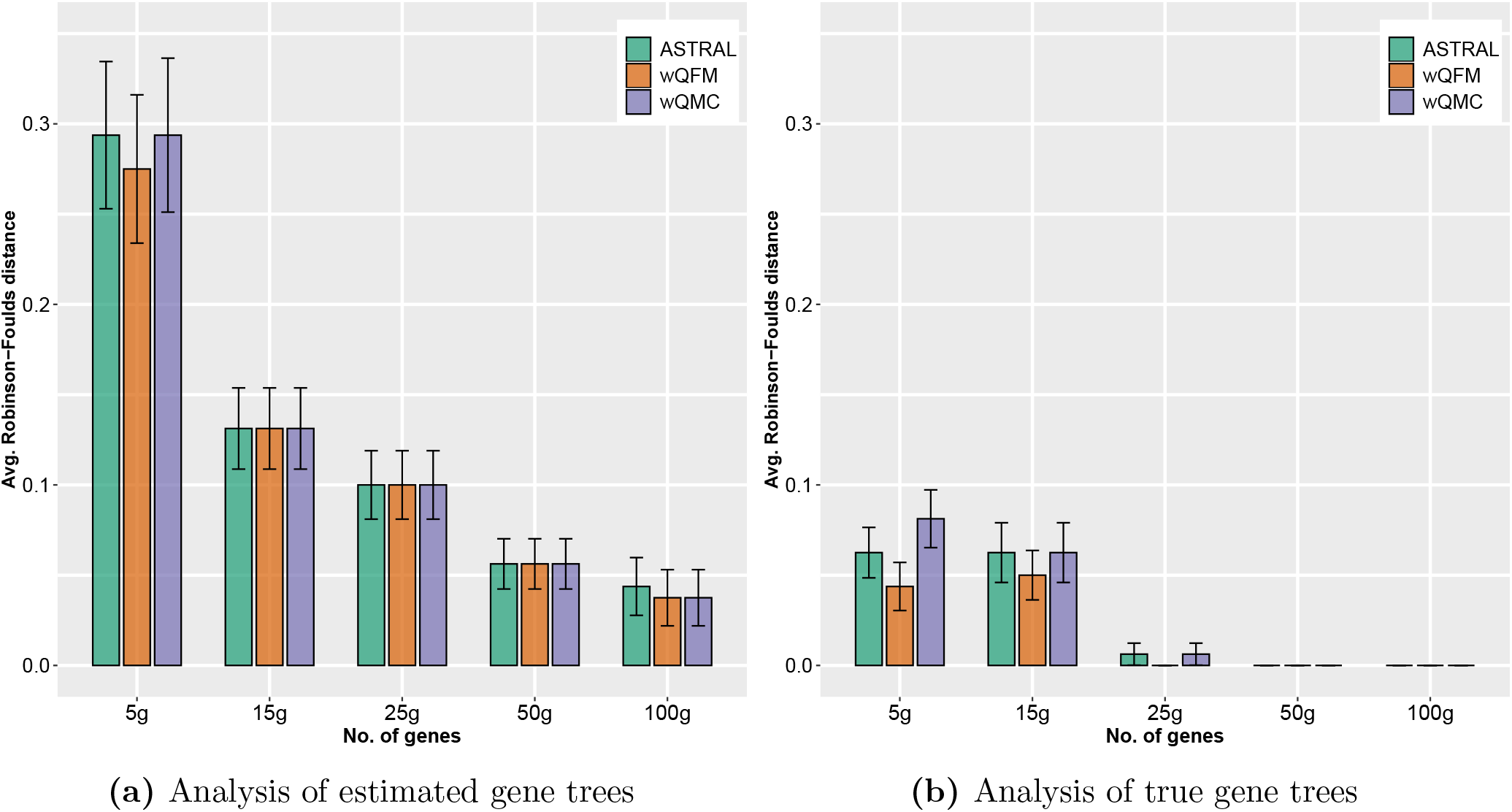
Comparison of ASTRAL, wQFM and wQMC on 11-taxon high-ILS dataset. We varied the number of genes (5 genes to 100 genes) for both estimated and true gene trees. We show the average RF rates with standard errors over 20 replicates.

On this smaller dataset, ASTRAL was run with its exact version, meaning that it is guaranteed to return a tree with the highest quartet score. Therefore, the fact that wQFM is sometimes more accurate than ASTRAL (albeit by a narrow margin) is interesting. These results support that, on real biological datasets with limited numbers of genes, summary methods tend to “overshoot” the optimization criteria that they try to optimize [65]. These results also support the presence of “phylogenomic terrace”, where multiple trees with the same quartet score can have different topological accuracies, and multiple trees with the same topological accuracy may have different quartet scores (see [65] for more details, and also [63]). Therefore, different algorithmic design for searching the species tree space under the quartet score may lead to different tree accuracies even though they achieve similar quartet scores.

### 4.3 Results on 48-taxon avian simulated dataset

Fig. 3(a) shows the performance on varying the ILS levels (0.5X, 1X, 2X) with 1000 genes and fixed default sequence length (500 bp). Unlike 37-taxon dataset, ASTRAL outperformed both wQFM and wQMC. wQFM was the second best method, outperforming wQMC on all three ILS levels, and the differences were statistically significant for 0.5X and 1X model conditions. We also assessed the performance on varying numbers of genes with fixed default level of ILS (1X level) and 500bp sequence length (Fig. 3(b)). wQFM matched the accuracy of ASTRAL (except for the 500- and 1000-gene model conditions), whereas wQMC incurred the highest tree errors. The improvement of wQFM and ASTRAL over wQMC are statistically significant in most cases.

**Figure 3:**
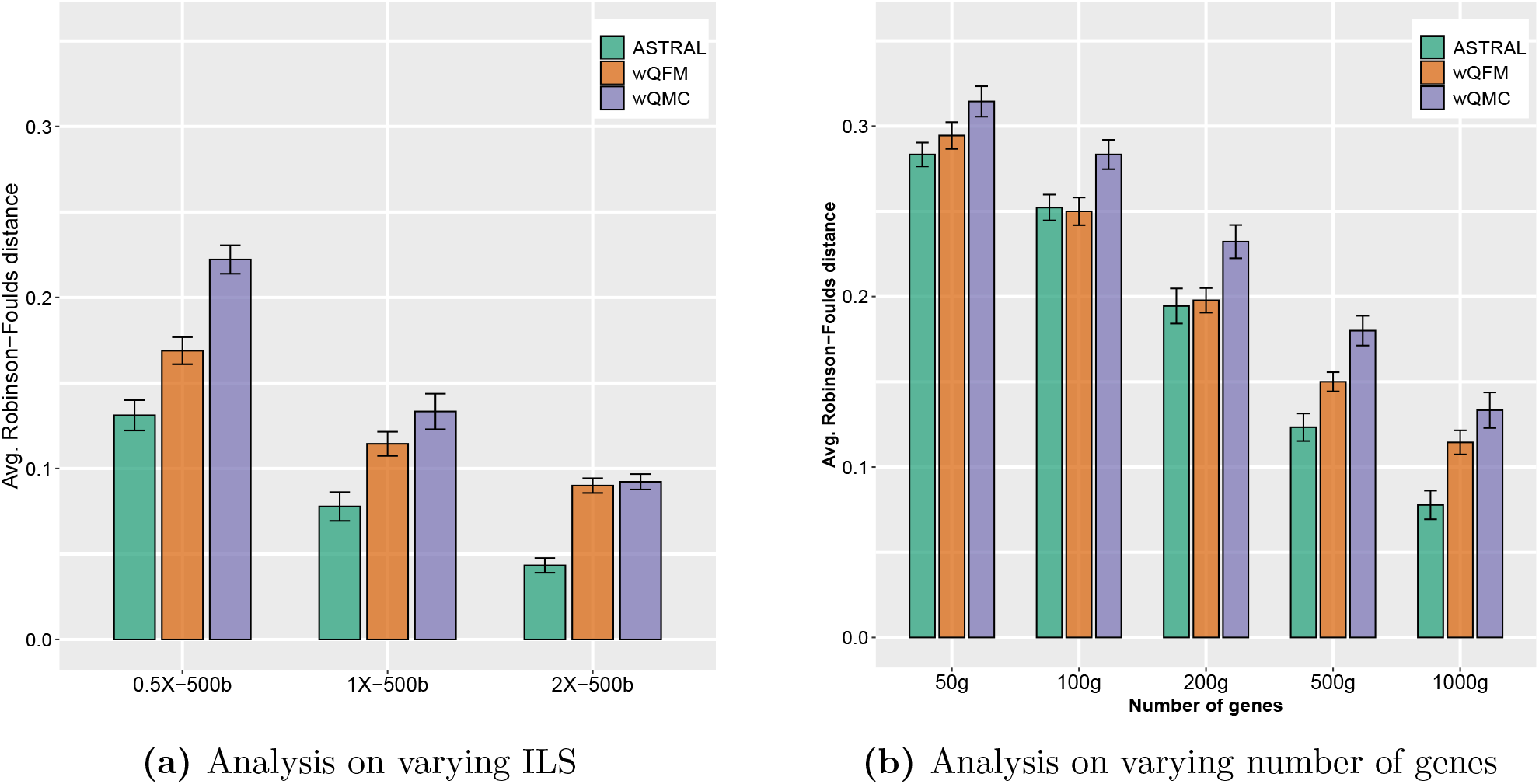
Comparison of ASTRAL, wQFM and wQMC on 48-taxon simulated avian dataset. We show the average RF rates with standard error bars over 20 replicates. (a) The level of ILS was varied from 0.5X (highest) to 2X (lowest) amount, keeping the sequence length fixed at 500bp and the number of genes at 1000, (b) number of genes was varied from 50g to 1000g, with 500bp sequence length and moderate (1X) ILS.

### 4.4 Results on 15-taxon and 101-taxon dataset

Similar trends were observed on 15-taxon and 101-taxon dataset (see Sec. 3 in the supplementary materials.).

### 4.5 Results on biological dataset

#### Avian Dataset

We have re-analyzed the avian biological dataset containing 14,446 loci across 48 taxa. This is an extremely challenging dataset since it contains high levels of gene tree discord, perhaps because their ancestors experienced a rapid radiation [51]. The original analyses of this data by MP-EST using the binned gene trees was largely congruent with combined analyses using ExaML [66], and both trees were presented as reference [51, 56].

Similar to the unbinned MP-EST analysis [51,56,67], ASTRAL, wQFM and wQMC run on 14,446 unbinned gene trees violate several subgroups established in the avian phylogenomics project and other studies (indicated in red in Fig. 4). However, wQFM estimated tree is more closer to the reference MP-EST tree (on binned gene trees) than the trees estimated by ASTRAL and wQMC. ASTRAL and wQMC differ in 9 and 10 edges, respectively, with respect to the reference MP-EST tree, whereas wQFM differs in 6 edges. Notably, the reference combined analysis tree and the MP-EST tree (presented in [51]) differ with each other in 5 edges.

**Figure 4:**
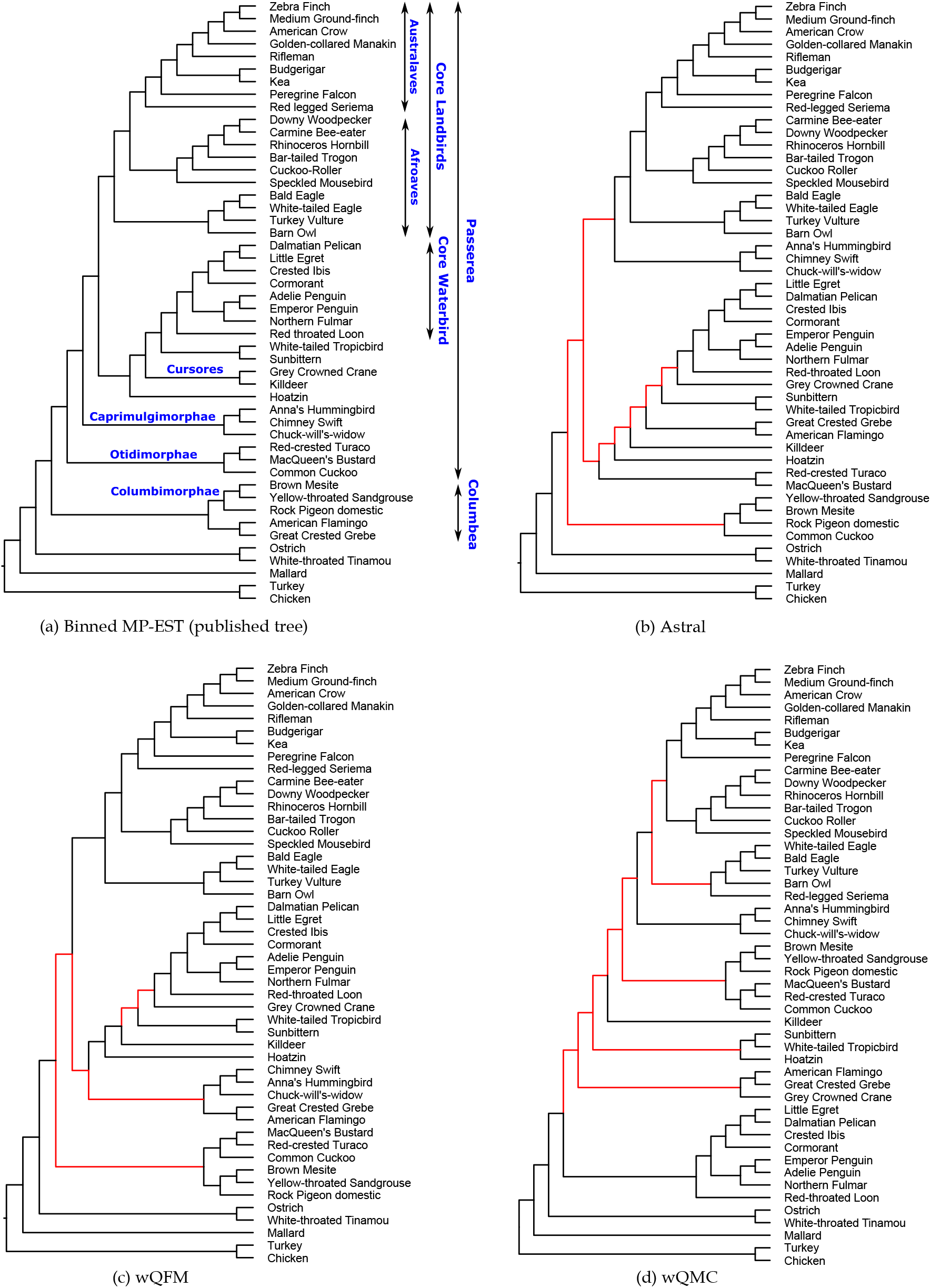
Estimated trees on the avian dataset with 14,446 genes. (a) Reference trees from the original paper [51] using MP-EST with statistical binning [56], (b)-(d) trees estimated by ASTRAL, wQFM and wQMC, respectively on 14,446 unbinned gene trees. Branches conflicting with the reference coalescent-based tree are shown in red lines.

Both wQFM and ASTRAL correctly placed seriemas as sister to the clade containing passerimorphae (passeriformes, parrot) and falcon, and thus reconstructed the well established Australaves clade (passeriformes, parrot, falcon, seriema). However, wQMC misplaced seriemas and failed to recover Australaves. ASTRAL and wQFM correctly reconstructed the core landbirds and core waterbirds [51,68–71]. wQMC also grouped them together, albeit with different branching orderings. wQFM correctly identified the core waterbirds as belonging to the sister clade of the core landbirds, whereas both ASTRAL and wQMC misplaced Caprimulgimorphae as sister to the core landbirds. While these methods recovered the core landbirds and core waterbirds [51,68–71] as well as various smaller sub-groups (e.g., Passerimorphae, Accipitrimorphae, Phaethontimorphae, Caprimulgimorphae, Phoenicopterimorphae), they could not recover some key clades. All of them failed to recover Columbea (flamingo, grebe, pigeon, mesite, sandgrouse). Although all of them correctly constructed Columbimorphae (mesite, sandgrouse, pigeon) and Phoenicopterimorphae (flamingo, grebe), they did not place them as sister clades and thus failed to recover Columbea. ASTRAL failed to recover Otidimorphae (bustard, turaco, cuckoo), whereas both wQFM and wQMC reconstructed this clade. All these methods failed to recover Cursores (crane, killdeer).

#### Mammalian Dataset

We re-analyzed the mammalian dataset from [59] containing 447 genes across 37 mammals after removing 21 mislabeled genes (confirmed by the authors), and two other outlier genes. The trees produced by wQFM, wQMC and Astral are identical to each other (see Fig. 5(a)). This tree placed tree shrews (*Tupaia belangeri*) as sister to Glires, which is consistent to the CA-ML analyses (reported in [59]), and bats have been placed as sister to the clade containing Cetartiodactyla, Carnivora, and Perissodactyla (which is consistent to the MP-EST analyses [72]). However, alternative relationships (e.g., tree shrew as sister to Glires, and bats as sister to Cetartiodactyla) have also been observed [15,72]. The placement of tree shrews and bats is of substantial debate [73–76], and so the differential placement is of considerable interest in mammalian systematics.

**Figure 5:**
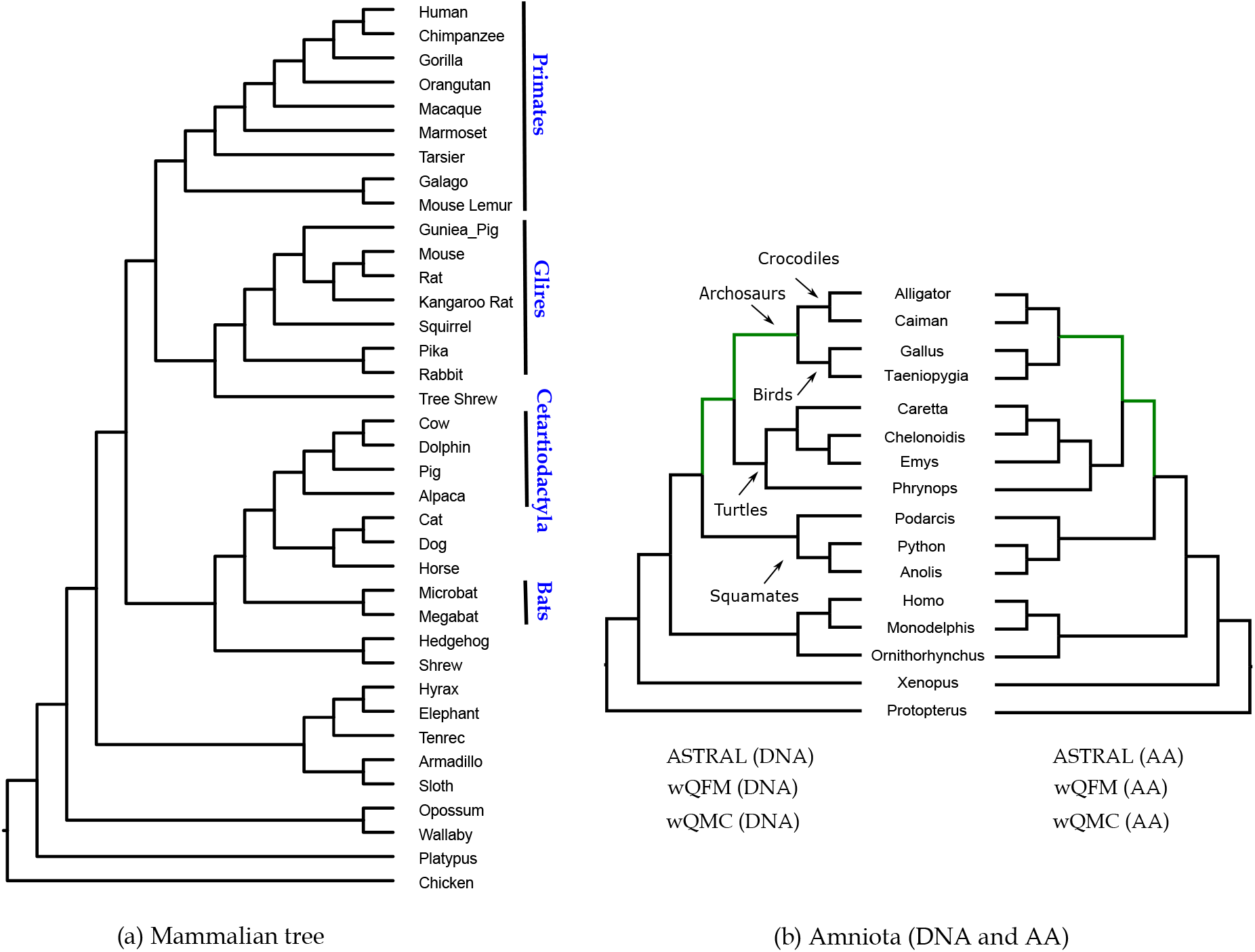
Analyses of the mammalian and amniota dataset using ASTRAL, wQFM and wQMC. (a) The tree estimated by ASTRAL, wQMC and wQFM on the mammlian dataset, (b) analysis of the amniota dataset using both DNA and AA gene trees.

#### Amniota dataset

We re-analyzed the amniota dataset (both amino acid (AA) and nucleotide (DNA) gene trees) from Chiari *et al*. [60] containing 248 genes across 16 amniota taxa. The goal is to resolve the position of turtles relative to birds and crocodiles. Previous studies [60,72,77,78] suggest the sister relationship between birds and crocodiles (forming archosaurs), and the placement of turtles as the sister to archosaurs.

ASTRAL, wQMC and wQFM, either on AA or DNA gene trees, correctly put turtles as a sister clade to archosaurs (see Fig. 5(b)). All these three methods, either on AA or DNA data, reconstructed identical trees and these two trees (on AA and DNA data) are highly congruent, differing only in the resolution of squamates (lizards and snakes).

#### Angiosperm dataset

We analysed the angriosperm dataset from Xi *et al*. [61] containing 310 genes samples from 42 angiosperms and 4 outgroups. The key question here is to investigate the position of *Amborella trichopoda Baill*. Our analyses with ASTRAL, wQFM and wQMC support the placement of *Amborella* as sister to water lilies (i.e., Nymphaeales) and rest of the angiosperms (see Fig. 6). This placement of *Amborella* is congruent to the CA-ML analysis in Xi *et al*. [61] and other molecular studies [58,79,80]. An alternate hypothesis, which supports a clade containing *Amborella* plus water lilies that is sister to all other angiosperms, has also been observed [61,81,82]. wQFM and wQMC differ from ASTRAL on a single edge (the placement of Sapindales (Citrus)).

**Figure 6:**
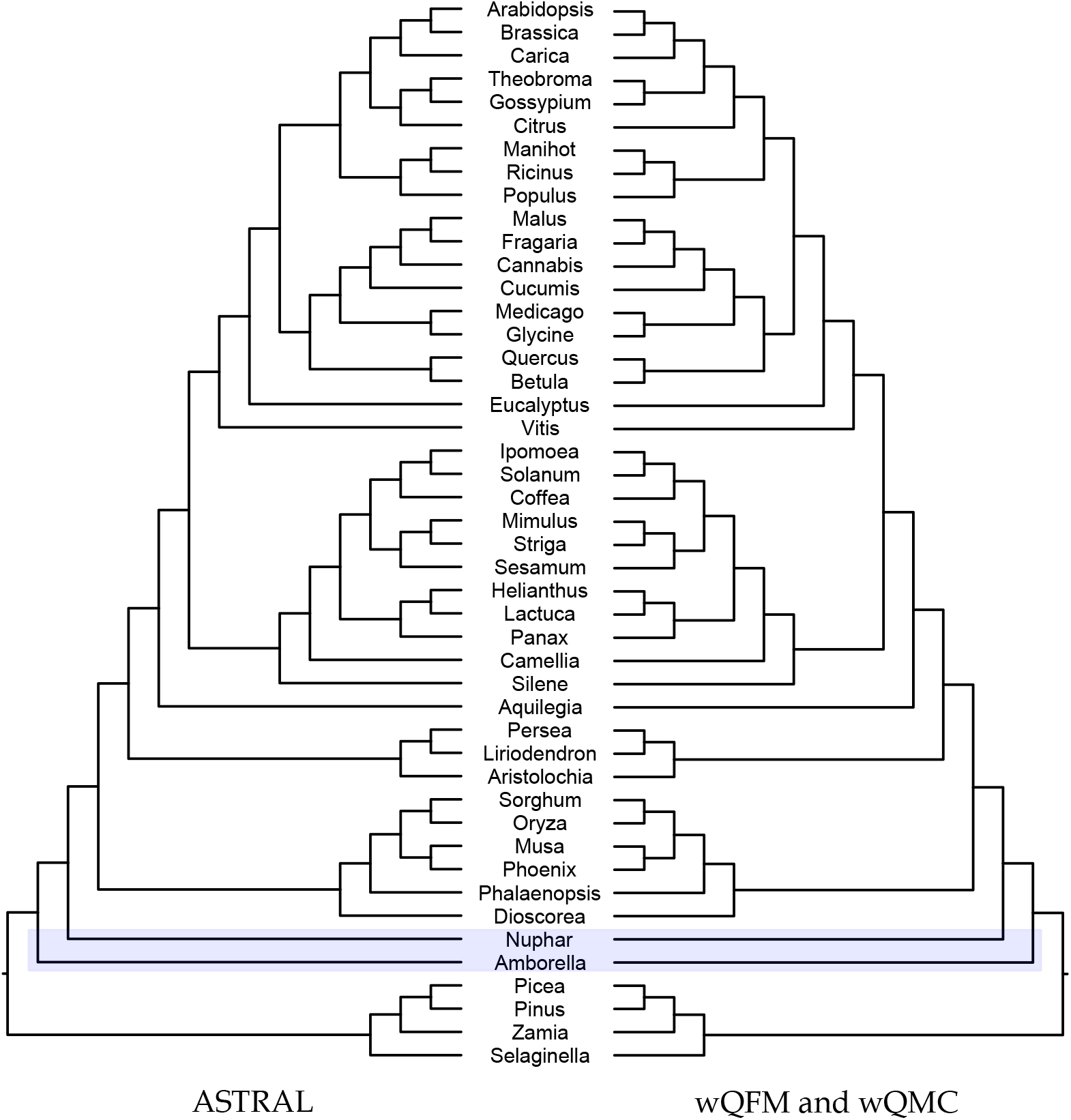
Analyses of the angiosperm dataset using ASTRAL, wQFM and wQMC. All these three methods support the placement of *Amborella* alone as sister to all other extant angiosperms.

### 4.6 Comparison with QFM

Both QFM and wQFM construct trees from a collection of quartets, but wQFM is capable of handling weighted quartets. One of the motivations for using the weighted version is to use all possible quartets (i.e., 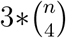 quartets for n taxa) with relative weights instead of using the unweighted setting, where – for each set of four taxa – we are given with one of the three alternate quartet topologies [20,24]. In order to show the efficacy of using weighted quartets, we have compared wQFM with QFM (see supplementary materials). Experimental results suggest that assigning weights can improve phylogenomic analysis.

**Table 1:**
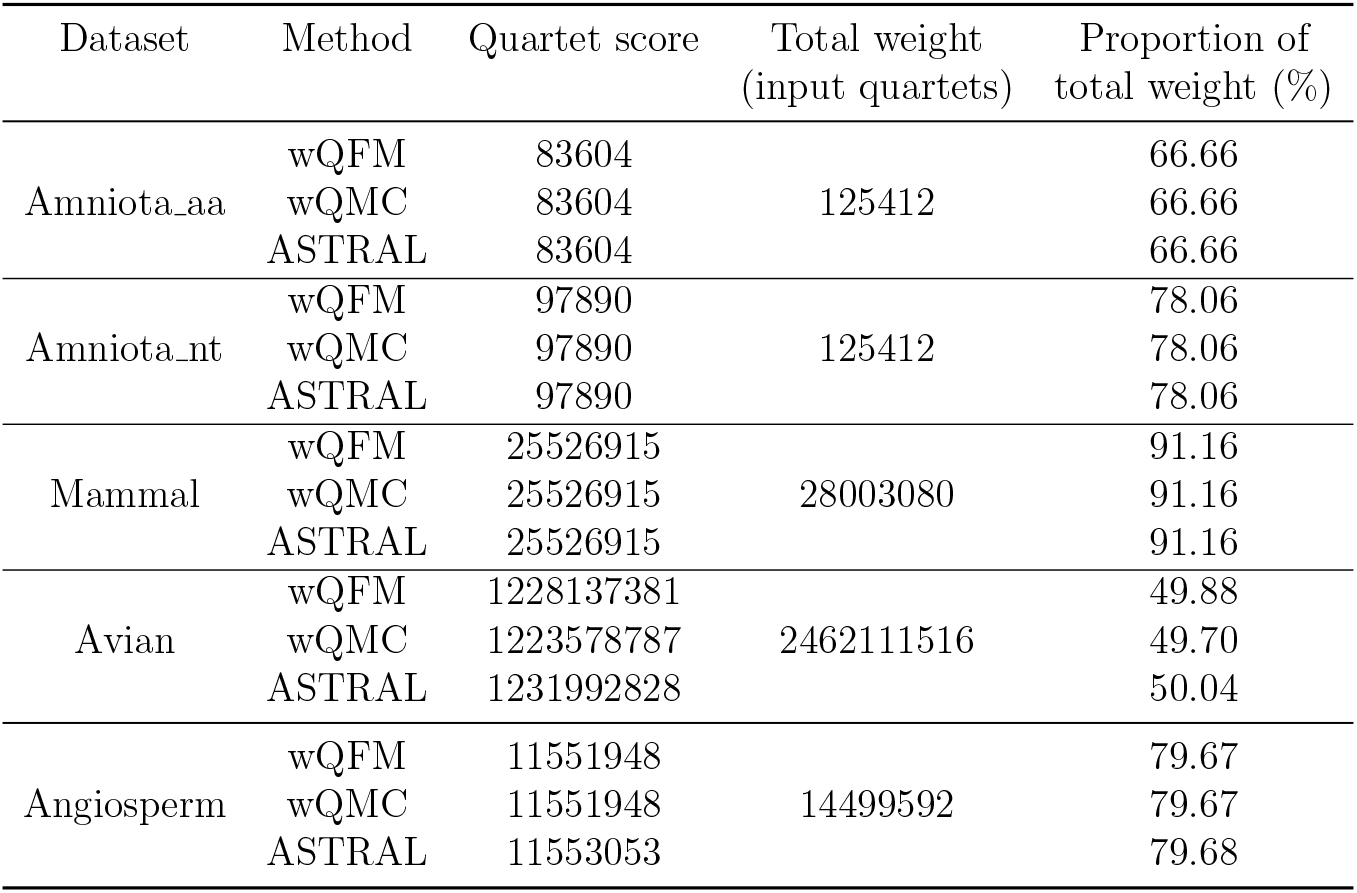
Quartet scores of various methods on biological datasets. We show the quartet scores (sum of the weights of the satisfied quartets) of various methods, total weight of the quartets in the input gene trees, and their respective ratios.

| Dataset | Method | Quartet score | Total weight<br>(input quartets) | Proportion of<br>total weight (%) |
| --- | --- | --- | --- | --- |
| Amniota_aa | wQFM | 83604 | 125412 | 66.66 |
|  | wQMC | 83604 |  | 66.66 |
|  | ASTRAL | 83604 |  | 66.66 |
| Amniota_nt | wQFM | 97890 | 125412 | 78.06 |
|  | wQMC | 97890 |  | 78.06 |
|  | ASTRAL | 97890 |  | 78.06 |
| Mammal | wQFM | 25526915 | 28003080 | 91.16 |
|  | wQMC | 25526915 |  | 91.16 |
|  | ASTRAL | 25526915 |  | 91.16 |
| Avian | wQFM | 1228137381 | 2462111516 | 49.88 |
|  | wQMC | 1223578787 |  | 49.70 |
|  | ASTRAL | 1231992828 |  | 50.04 |
| Angiosperm | wQFM | 11551948 | 14499592 | 79.67 |
|  | wQMC | 11551948 |  | 79.67 |
|  | ASTRAL | 11553053 |  | 79.68 |

### 4.7 Running time

We performed the experiments on a Linux machine with 8 GB RAM and i7 2.50 GHz processor. We ran the exact version of ASTRAL-III (version: 5.7.3) for smaller datasets (11 ~ 15 taxa), and used the heuristic version to analyze larger datasets. For wQFM and wQMC, we report the running time for amalgamating the weighted quartets (given as input), which excludes the time for computing the weighted quartets. We used a custom script for generating weighted quartets, by computing the frequency of each quartet, from a collection of gene trees. However, weight/confidence of a quartet can be generated in different ways, e.g., using the quartet frequency (as used in this study), likelihood of a quartet, and various algebraic and statistical approaches. Thus, the time required to generate weighted quartets may differ depending on what types of weights are being used.

For smaller datasets, these three methods wQFM, wQMC and ASTRAL took very small amounts of time. Both wQFM and wQMC took only a fraction of a second for 11 and 15-taxon datasets. ASTRAL also took around a second to analyze 11-taxon datasets. For 15-taxon datasets however, it took ASTRAL (exact version) 1 to ~12 min (depending on various numbers of genes), which is much longer than wQFM and wQMC. Note that, since the input to wQFM and wQMC are weighted quartets embedded in the input gene trees, their running times are not much sensitive to the number of genes.

For 37-taxon dataset, heuristic version of ASTRAL was used, which led to much smaller running times per replicates, ranging from 2 ~6 s. The running time of ASTRAL decrease from 6 s to 2 s as we decrease the level of ILS from higher ILS (0.5X) to lower ILS (2X). This decrease in running time may be due to the fact that as the amount of discordance (due to ILS) decreases in the gene trees, the number of bipartitions in the gene trees also decrease. This leads to a smaller search space for ASTRAL (heuristic version) to explore. ASTRAL took around 1 s (on 50 genes) to 10 s (on 800 genes) to analyze various numbers of genes. wQMC was the fastest method which took only a second to analyze a single replicate of this dataset. wQFM was also fast, taking only 4 ~12 s.

On the 48-taxon simulated avian dataset, wQMC took around 2 s. ASTRAL’s running time ranges from 6 ~150 s, depending on various numbers of genes and ILS levels. wQFM took around 15 ~35 s per replicate on avian dataset. The most significant difference in running times was observed on the avian biological dataset with 14K gene trees, where ASTRAL took ~32 hours to run. wQMC and wQFM, on the other hand, finished within 2 and 20 s, respectively. This is due to the fact that ASTRAL’s running time increases as we increase the number of genes, but wQMC and wQFM takes as input a set of weighted quartets and thus their running times are not sensitive to the number of genes. This is attributed to the fact that ASTRAL’s running time is much more sensitive to the number of genes (hence to the number of unique bipartitions) than wQFM and wQMC. On a relatively larger dataset with 101 taxa, the running time of wQFM ranges from 25 ~35 min. ASTRAL and wQMC were much faster, taking around 2 ~ 3 min and 5 s, respectively.

## 5 Conclusions

We present wQFM – a new statistically consistent method for estimating species trees from genome-scale data by amalgamating weighted quartets, which matches or improves upon the best existing methods under a range of realistic model conditions. Quartet amalgamation is an important class of methods which takes individual quartets as input and amalgamate them into a single coherent tree. With the recent advances in computing accurate quartet estimation using site pattern probabilities without needing to estimate gene trees [20], and thereby reducing the impact of gene tree estimation error in species tree estimation, quartet amalgamation techniques like QFM are being widely used and have drawn substantial attention from the systematists. Moreover, assigning relative weights to the quartets can potentially improve the tree accuracy [30], and thus can be more applicable than weight-oblivious techniques like QMC and QFM for estimating species trees from multi-locus data. wQMC and ASTRAL are two well known quartet based methods that can take the weights of the quartets into account. As previous studies have suggested the presence of phylogenomic terraces, i,e., multiple trees with an identical quartet score may have different topological accuracies (and vice versa) [65], developing various methods to efficiently search the tree space under the weighted quartet score will add value to the existing literature.

wQFM is a new divide-and-conquer based method for amalgamating weighted quartets. Our wide-ranging experimental results suggest that wQFM can reliably estimate species trees under practical and challenging model conditions. We showed that wQFM can estimate trees with similar or better accuracy than the main two alternate methods, wQMC and ASTRAL. Moreover, we showed that designing appropriate approaches for assessing a particular partition in the divide steps of divide-and-conquer based quartet amalgamation methods is crucial for computing reliable trees. We have proposed a novel scheme for finding partition scores which dynamically changes depending on the distribution of the weights of the quartets considered in a particular divide step. Thus, wQFM advances the state-of-the-art in weighted quartet amalgamation and can be used to accurately analyze upcoming multi-gene datasets. However, this study can be extended in several directions. This study is limited to small to moderate size datasets. Future studies need to investigate the performance of wQFM on relatively larger phylogenomic datasets with hundreds of taxa. This study considers the frequency of the quartets in input gene trees as weights. Future studies will need to investigate other ways of inferring quartet support including various statistical methods (e.g., likelihood score of a quartet with respect to sequence alignments) and their impact on the resulting trees. Both wQFM and wQMC consider the quartets present in the input set of gene trees. However, ASTRAL augments the set of bipartitions [62] in the gene trees and thus considers a larger tree space, which may result in better accuracies on large dataset. Exploring the performance of wQFM and wQMC on augmented sets of quartets by exploring the neighborhood of the input gene trees through SPR (subtree pruning and regrafting), NNI (nearest neighbor interchange) and TBR (tree bisection-reconnection) operations would be another interesting research direction. As we have shown that divide-and-conquer based quartet amalgamation methods could be sensitive to the way of computing partition scores, further investigations are required to design more effective approach for defining a partition score. wQFM is provably statistically consistent under the MSC model and has been evaluated on various datasets simulated under ILS. However, evaluating the performance of wQFM (weighted quartet amalgamation techniques in general) in the presence of horizontal gene transfer and gene duplications and losses would be interesting. On an ending note, our paper shows that the idea of estimating species trees by amalgamating weighted quartets has merit and should be pursued and used in the future phylogenomic studies.

## Supplementary Material

### 1 Overview

These supplementary materials present additional results, additional details on computing partition scores and the impact of various partition scores on the tree accuracy. Additionally, we observe the positive impact of including weights in both the partition scores as well as overall computation.

### 2 Effectiveness of using weighted quartets: comparison between wQFM and QFM

A direct comparison between wQFM and QFM using all possible quartets (weighted quartets for wQFM and unweighted quartet for QFM) sampled from the input set of gene trees is not meaningful. This is because using all possible 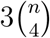 quartets *without any* weight as input and seeking a tree by maximizing the number of consistent quartets will just produce a random tree. Therefore, when using the unweighted setting – for each set of four taxa – the “best” quartets out of the three alternate quartet topologies are used [1]. In this study, we consider the support/weight for quartet tree *ab|cd* to be the number of the trees in 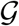 that induce *ab|cd* on set *a,b,c,d*. Therefore, for a set of four taxa *a,b,c,d*, the *best* quartet (out of three possible quartets: *ab|cd, ac|bd*, and *bc|ad*) is defined to be the quartet with the highest weight. In order to show the efficacy of the weighted setting, we compare the following three variants of quartet amalgamation.

1. wQFM with all possible weighted quartets.
2. wQFM with weighted best quartets. That means, 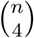 best weighted quartets (one quartet for each set of four taxa) are being used instead of 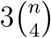 possible weighted quartets.
3. QFM with unweighted best quartets (one quartet for each set of four taxa).

Figures S1-S3 show the comparison among these three variants on various simulated datasets. These results show the superiority of weighted setting over the unweighted setting, as in most of the cases, wQFM with all possible weighted quartets outperformed the other two variants. Moreover, wQFM with weighted best quartets outperformed QFM with unweighted best quartets in many of the model conditions on these datasets – another evidence that assigning weights to the quartets can improve phylogenetic analyses.

**Figure S1:**
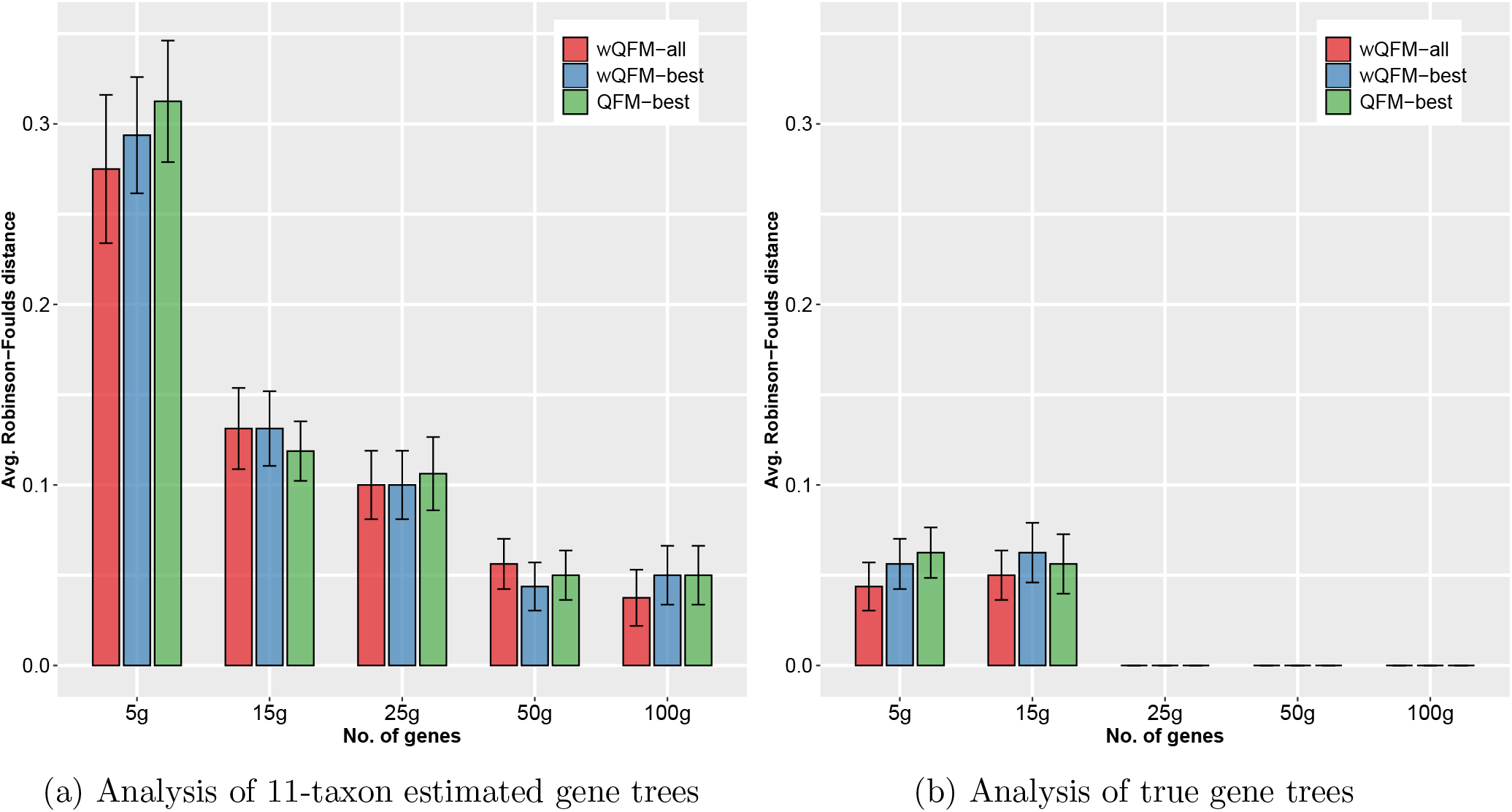
Comparison of wQFM and QFM on 11-taxon high-ILS dataset. We show the average RF rates over 20 replicates.

**Figure S2:**
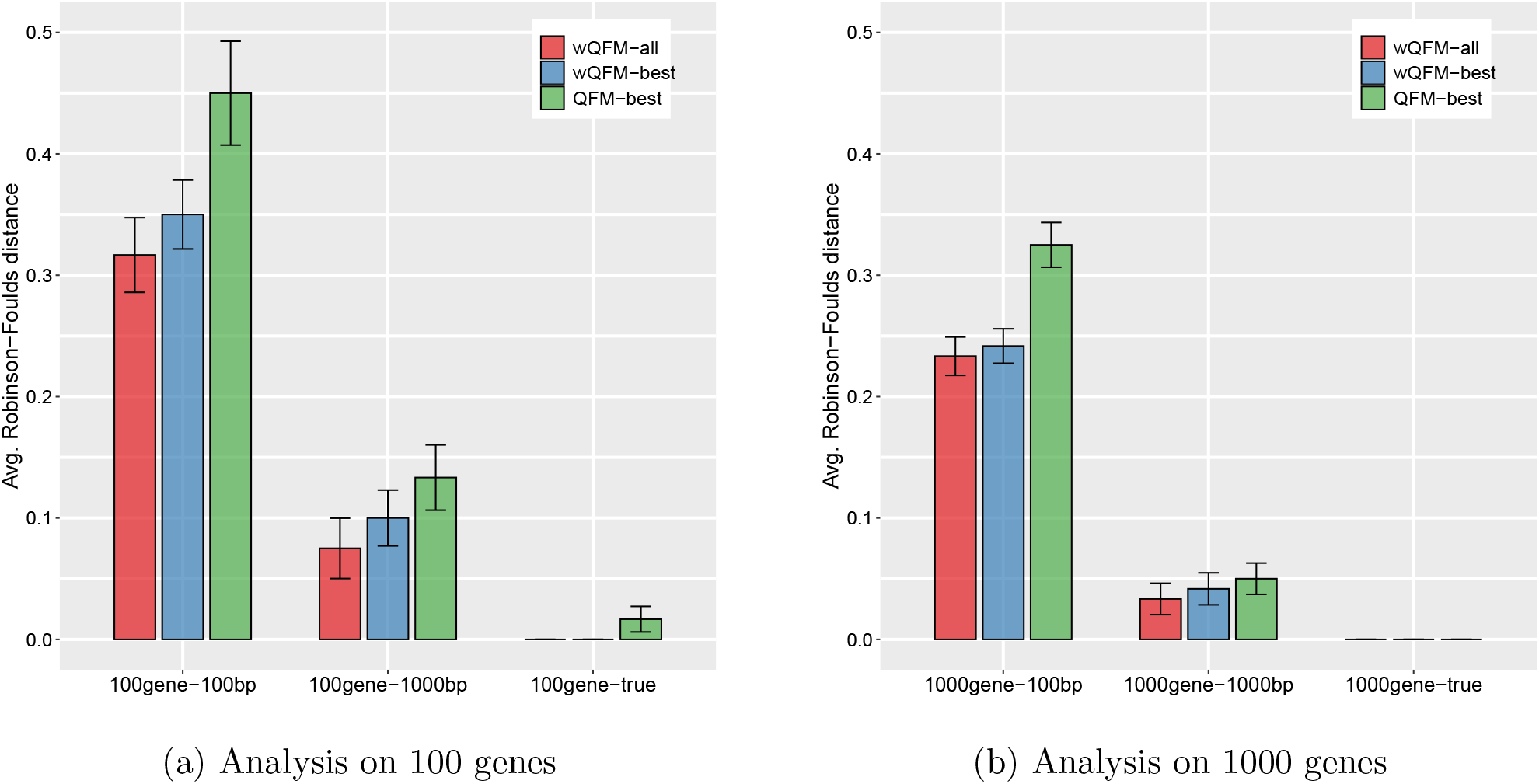
Comparison of wQFM and QFM on 15-taxon dataset. We show the average RF rates over 10 replicates.

**Figure S3:**
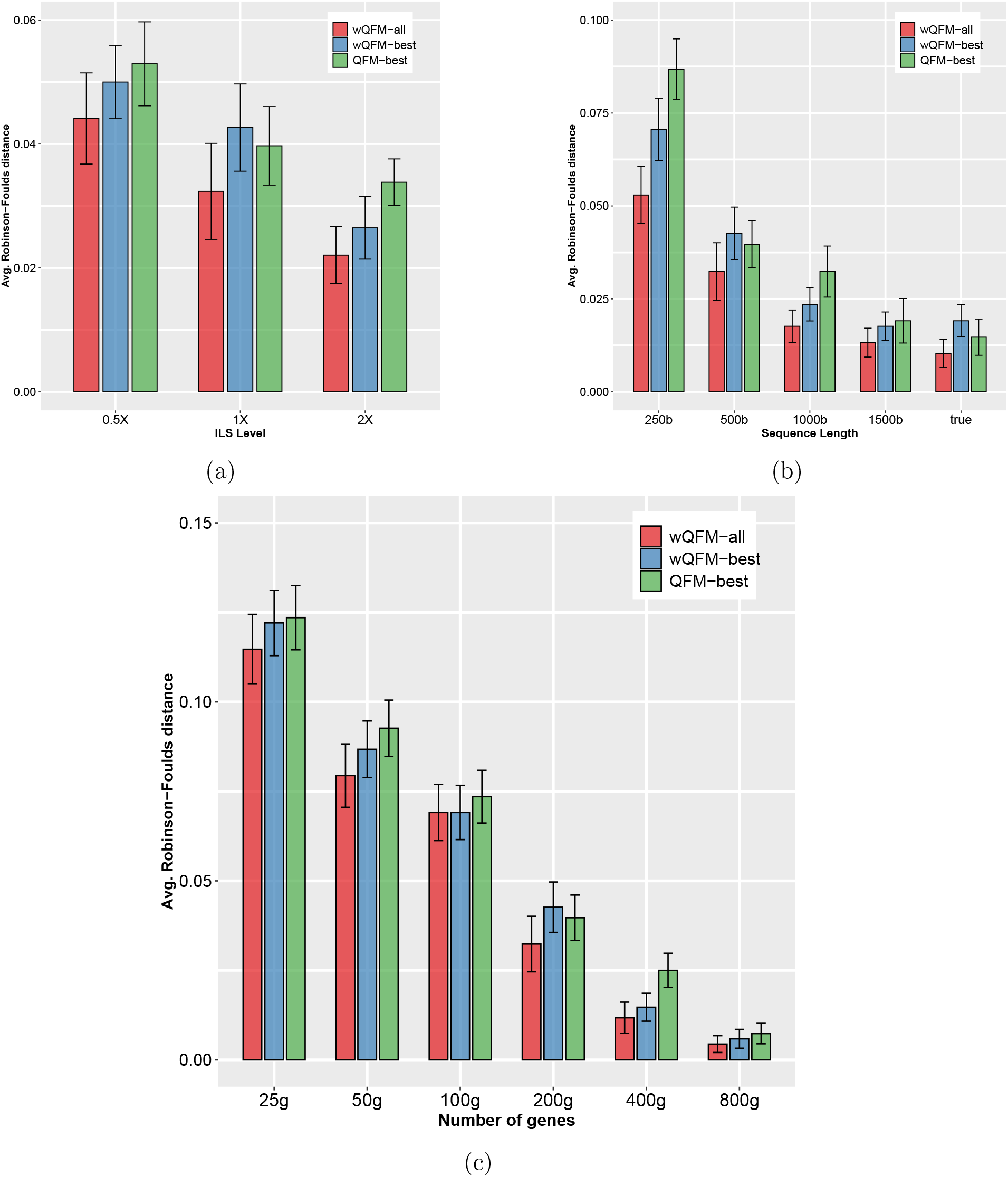
Comparison of wQFM and QFM on 37-taxon simulated dataset over 20 replicates per model condition. (a) The level of ILS was varied from 0.5X (highest) to 2X (lowest) amount, keeping the sequence length fixed at 500bp and the number of genes at 200; (b) The sequence length was varied from 250bp to 1500bp, keeping the number of genes fixed at 200, and ILS at 1X (moderate ILS); (c) The number of genes was varied from 25g to 800g, with 500bp sequence length and moderate (1X) ILS.

### 3 Algorithms for Computing Partition Scores

#### Algorithm 1

Dynamic Partition score calculation on each level

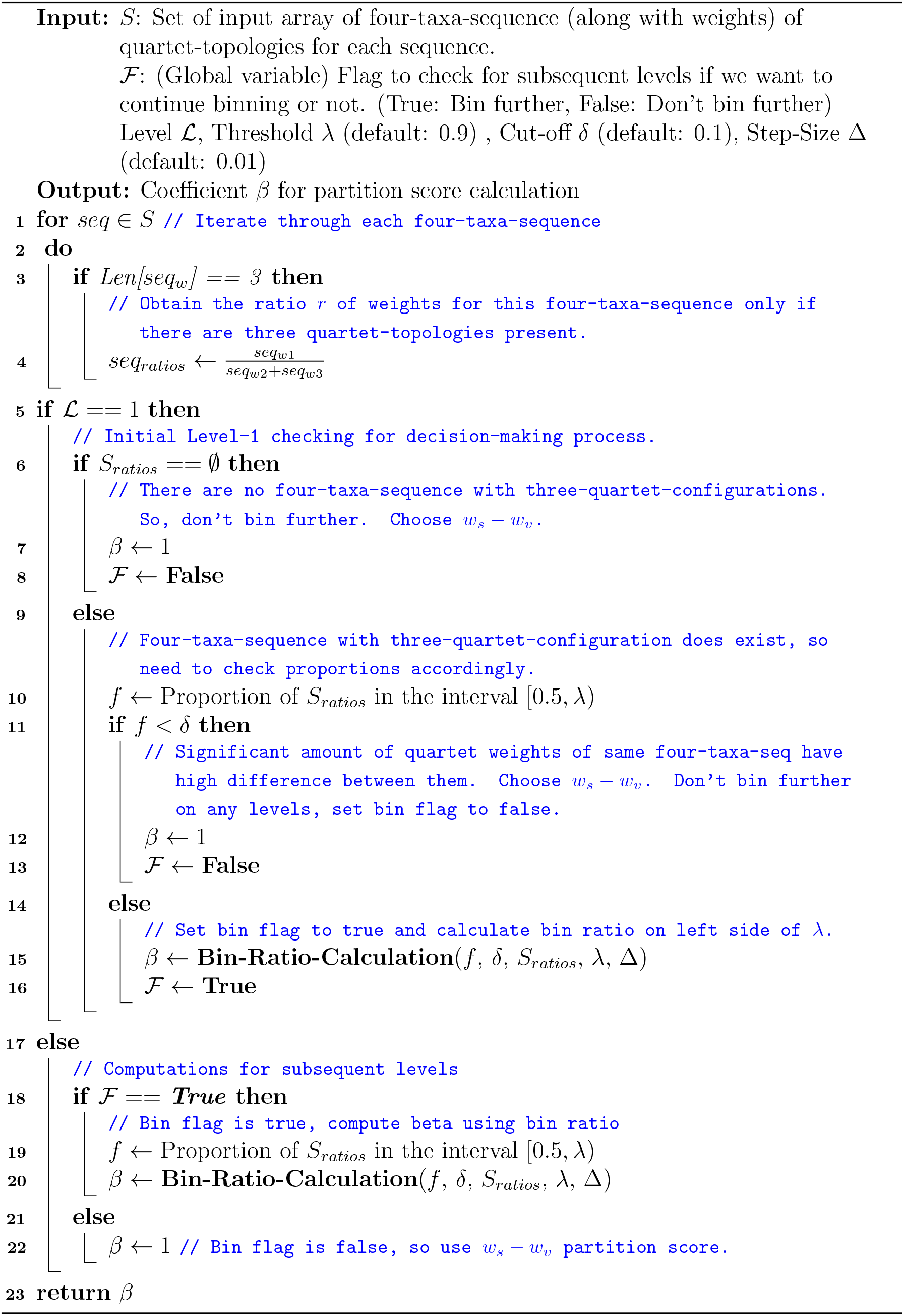

#### Algorithm 2

Function for Bin-Ratio-Calculation

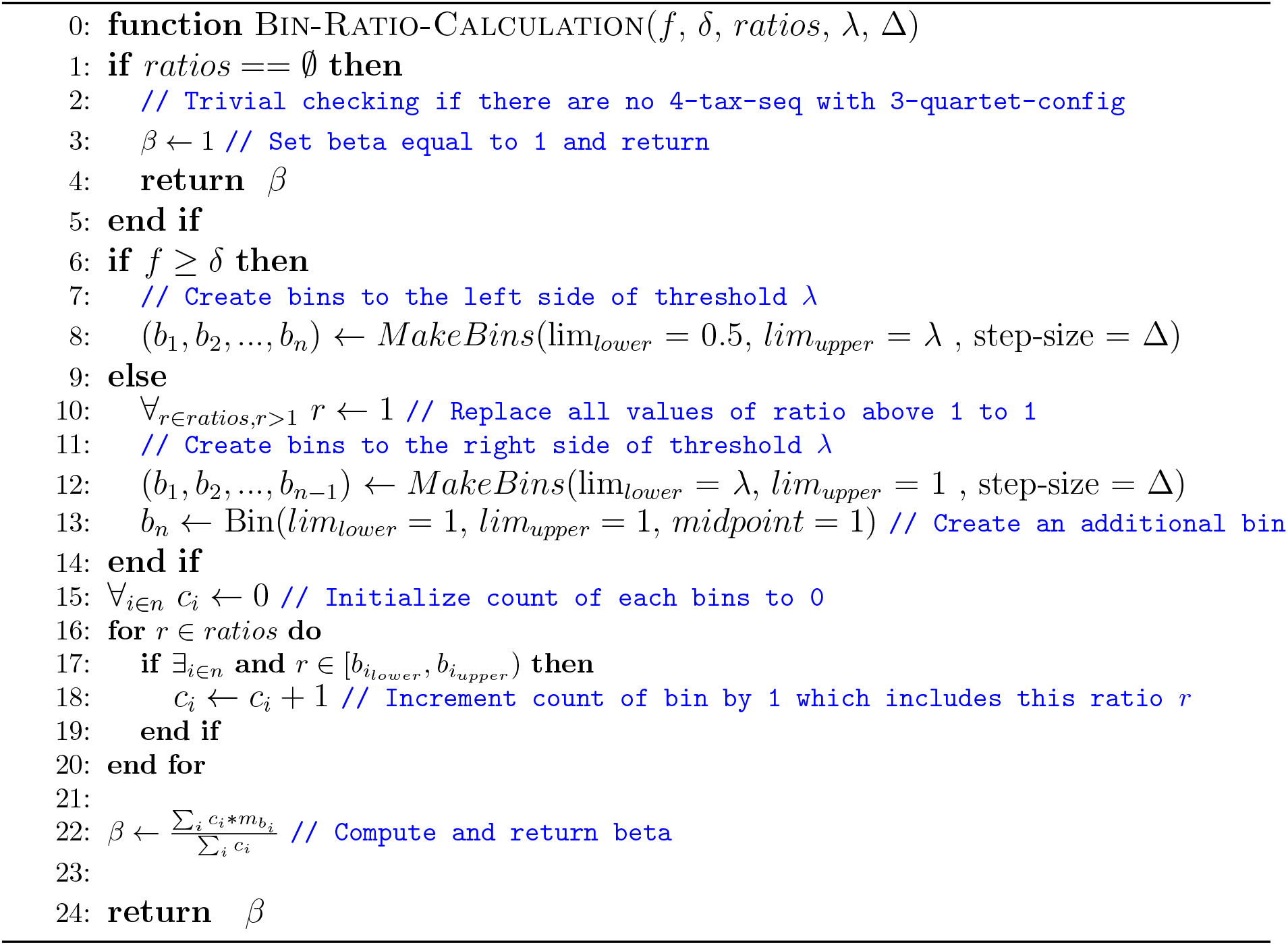

### 4 Additional Results

#### 4.1 Results on 15-taxon simulated dataset

We have explored the performance on varying gene tree estimation errors using 100bp and 1000bp sequence lengths, and numbers of genes (100 and 1000) as shown in Figure S4. ASTRAL, wQFM and wQMC achieved similar tree accuracy (with no statistically significant differences) on all the model conditions. All these methods improved as the number of genes and sequence length increased, and obtained best results (RF rate = 0) on true gene trees.

**Figure S4:**
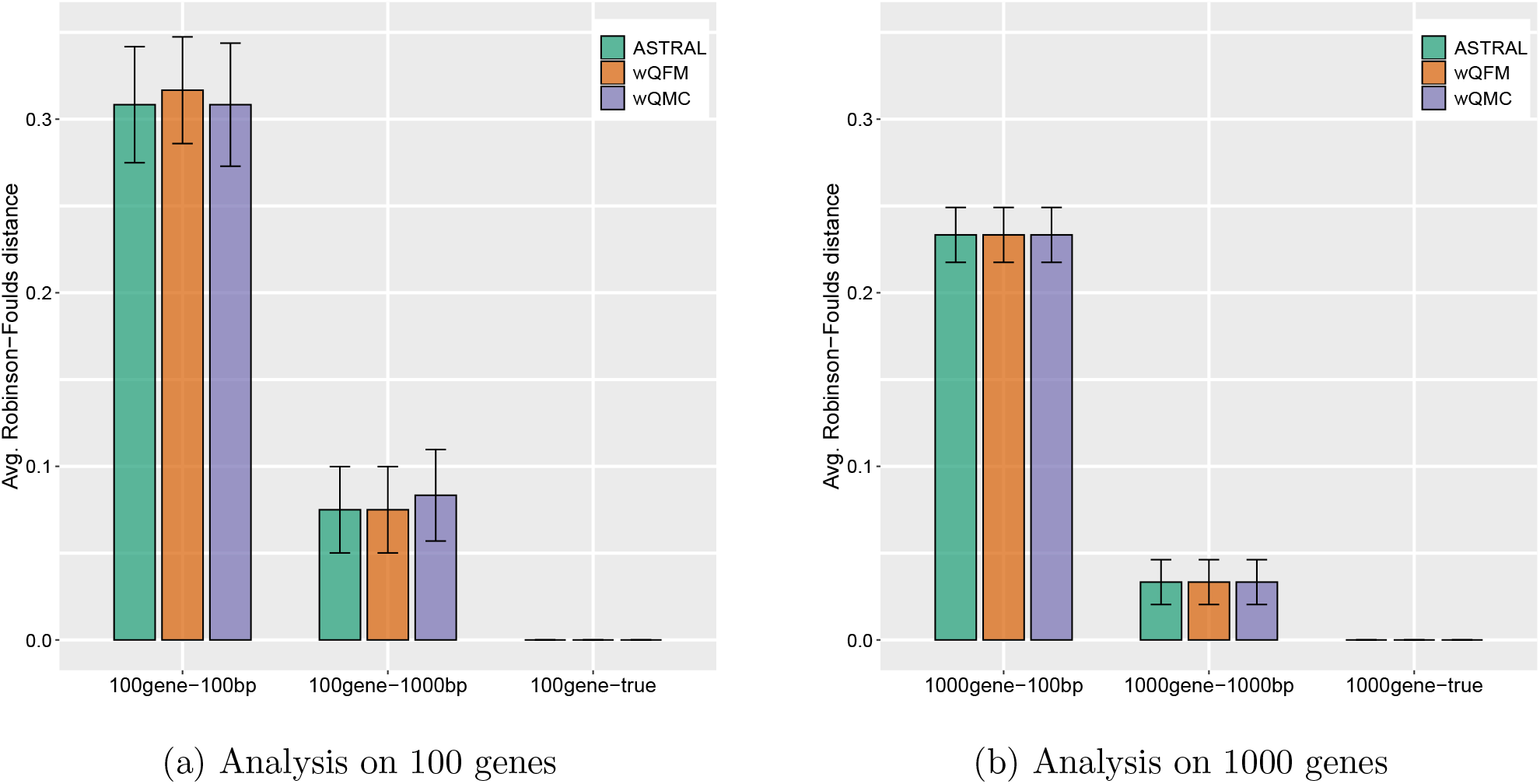
Comparison of ASTRAL, wQFM and wQMC on 15-taxon dataset. We show the average RF rates with standard errors over 10 replicates.

#### 4.2 Results on 101-taxon simulated dataset

We show average RF rates of wQFM, ASTRAL and wQMC on 10 replicates of 101-taxon dataset with 1000 true gene trees (see Fig. S5). All of these methods produced highly accurate trees with around 1.2% ~2.5% tree error. ASTRAL was the most accurate method followed by wQFM. However, the difference between them was very small and was not statistically significant.

**Figure S5:**
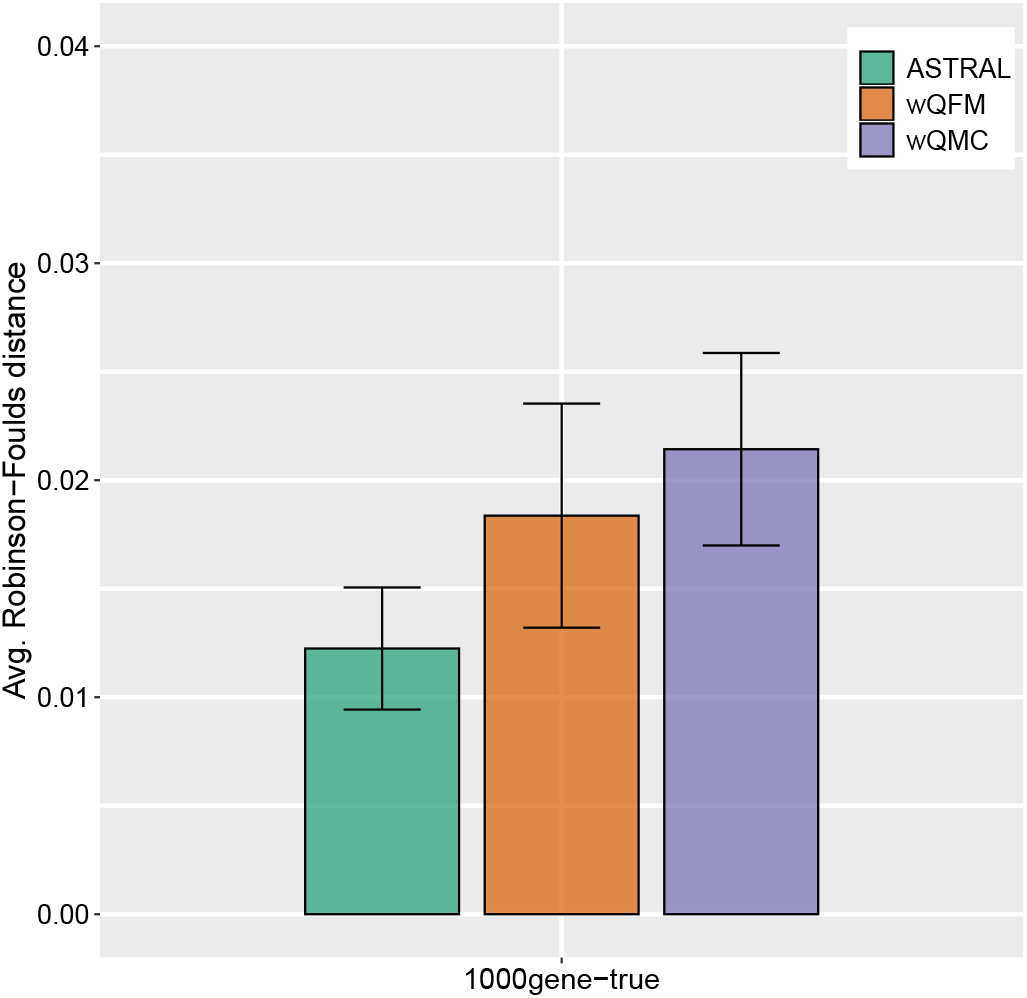
Comparison of ASTRAL, wQFM and wQMC on 101-taxon dataset. We show average RF rates with standard error bars over 10 replicates.

#### 4.3 Impact of partition scores on tree accuracy

The quality of the bipartitions generated by divide-and-conquer based methods like wQFM depend on the way a partition score is defined. We investigated how the tree accuracy of wQFM is affected by various types of partition scores. We considered the following four different partition scores, and evaluated wQFM, when it is run with these partition scores, on various simulated datasets (see Figs. S6, S7, S8, S9). Two of these four partition scores are *fixed*, meaning that we used a particular partition score in every divide steps of wQFM, and two of them are *dynamic*, meaning that the partition scores are not fixed, rather they are defined based on the distribution of the weights of the quartets (as described in Sec. 2.3 of the main paper).

- Fixed partition scores

1. *w_s_ − w_υ_*
2. *w_s_* − 0.5 ∗ *w_υ_*
- Dynamic partition scores

1. Dynamic Level 1: partition score is defined based on the distribution of the weights in the very first divide step, and the same score is used for subsequent divide steps.
2. Dynamic All Levels: partition scores are defined in each divide step based on the distribution of the weights of the quartets, which are being considered in a particular divide step. The results presented in this paper are based on this scheme.

**Figure S6:**
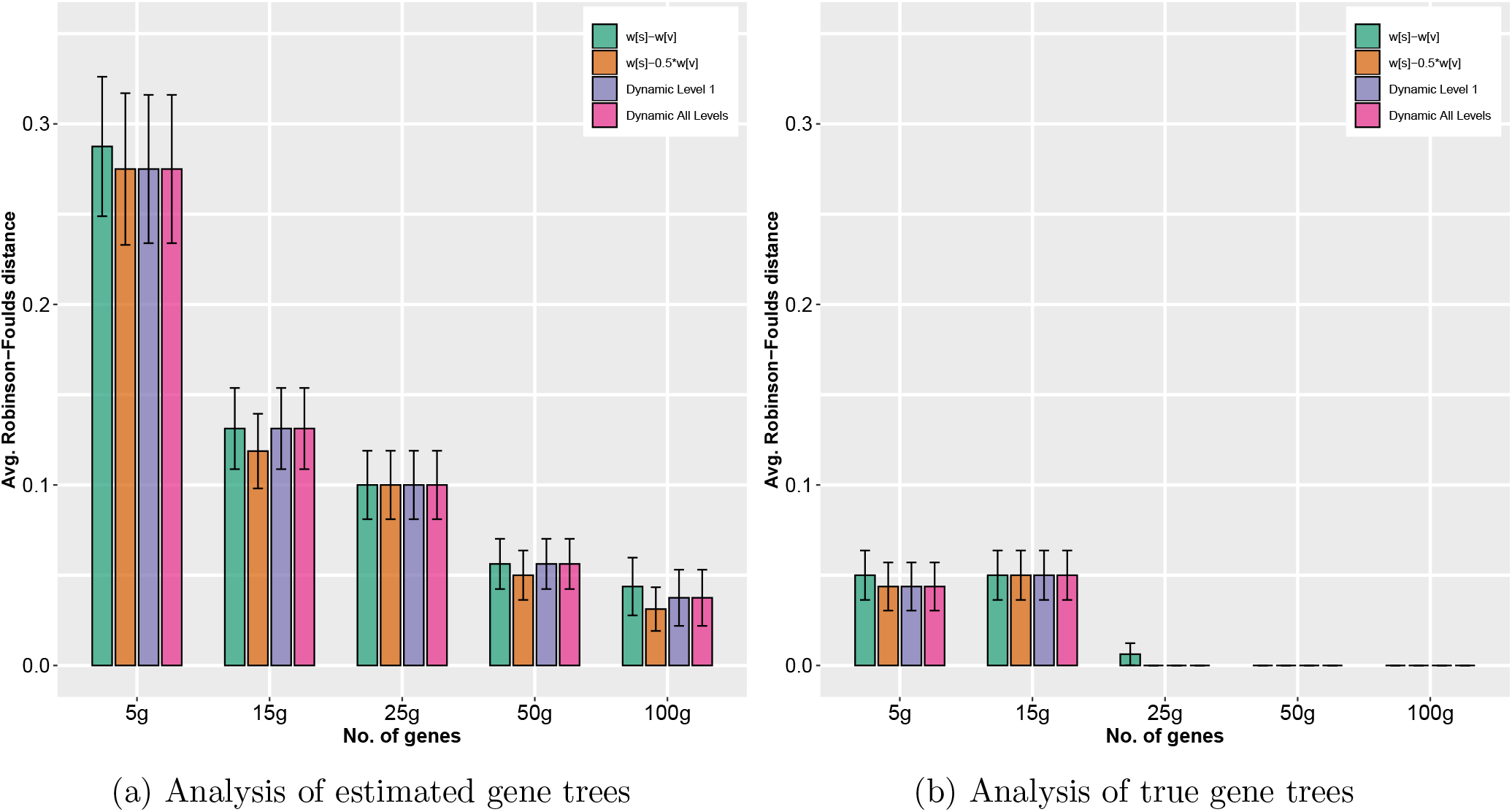
Impact of various partition scores on the accuracy of wQFM. We show average RF rates with standard error bars (over 20 replicates) on 11-taxon dataset by varying genes from 5g to 100g.

**Figure S7:**
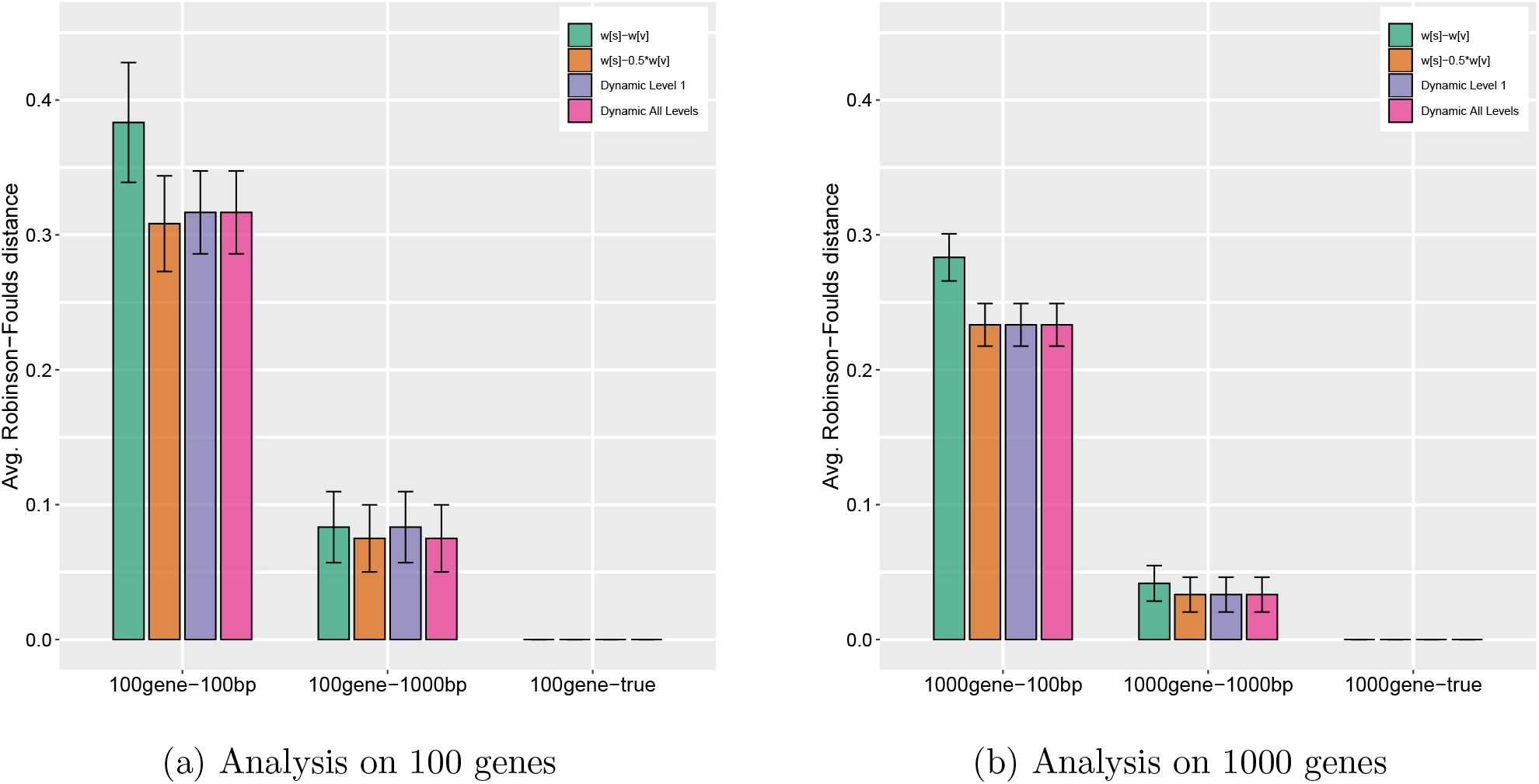
Impact of various partition scores on the accuracy of wQFM. We show average RF rates with standard error bars (over 10 replicates) on 15-taxon dataset.

**Figure S8:**
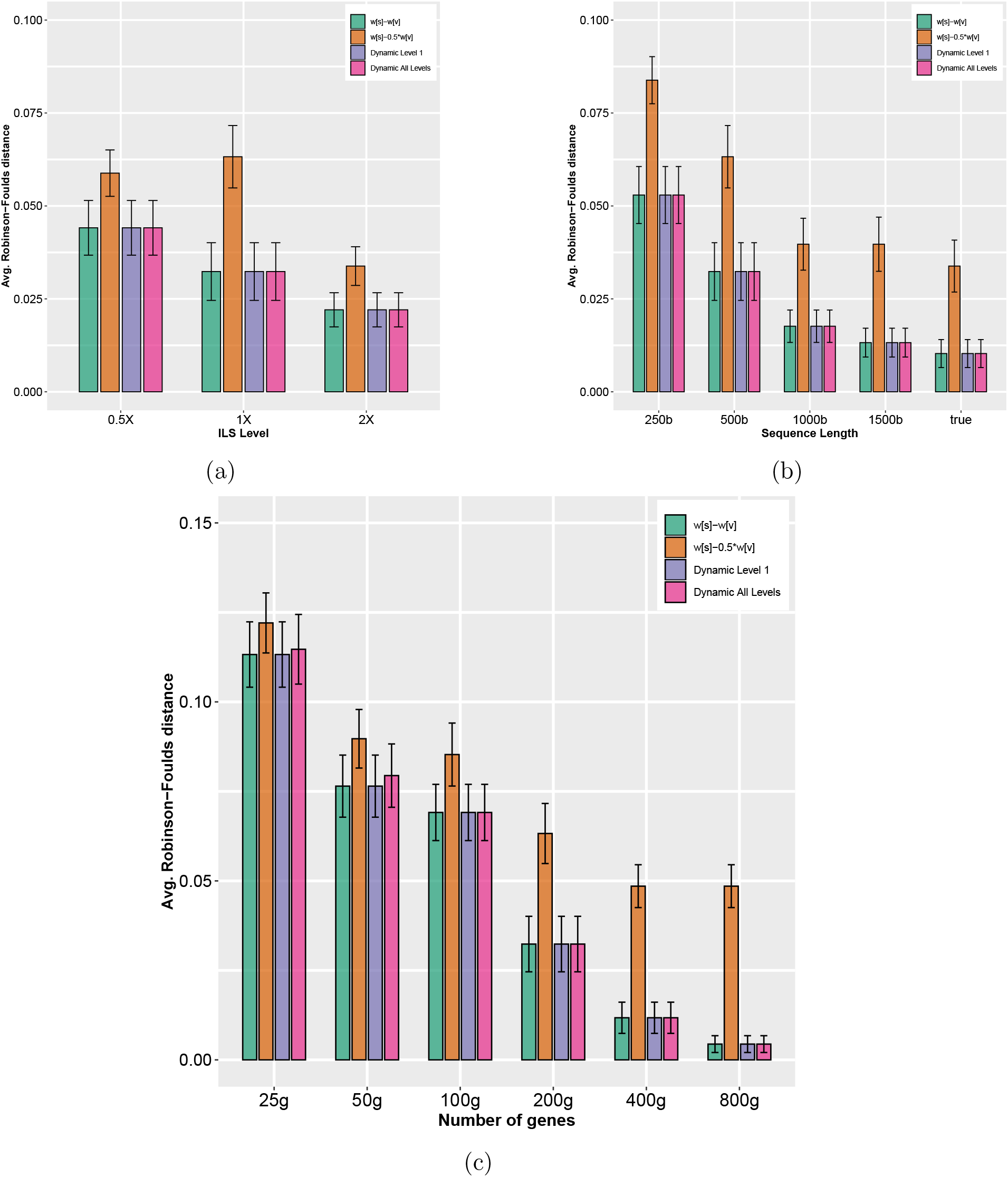
Impact of various partition scores on the accuracy of wQFM in 37-taxon dataset. We show average RF rates with standard error bars (over 20 replicates). (a) The level of ILS was varied from 0.5X (highest) to 2X (lowest) amount, keeping the sequence length fixed at 500bp and the number of genes at 200; (b) The sequence length was varied from 250bp to 1500bp, keeping the number of genes fixed at 200, and ILS at 1X (moderate ILS); (c) The number of genes was varied from 25g to 800g, with 500bp sequence length and moderate (1X) ILS.

**Figure S9:**
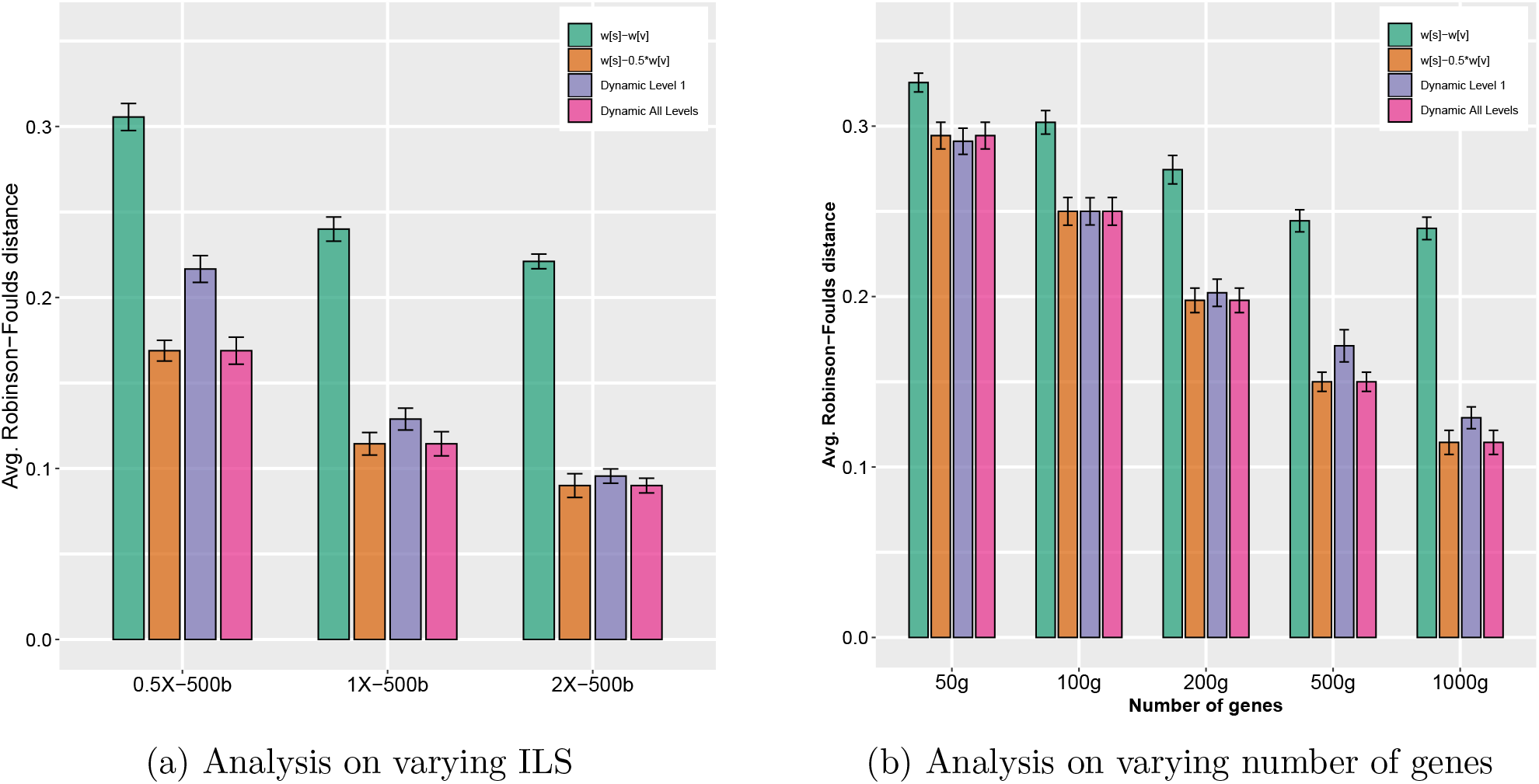
Impact of various partition scores on the accuracy of wQFM in 48-taxon dataset. We show average RF rates with standard error bars (over 20 replicates).

#### 4.4 Quartet Scores

Tables S2, S1, S3, and S4 show the quartet scores of various methods on various simulated datasets (11-, 15-, 37- and 48-taxon). The average absolute quartet-scores along with the average total weight of weighted quartets, and the average normalized quartets score per model condition is shown for the methods wQFM, wQMC, ASTRAL-III.

**Table S1:** Quartet scores on 15-taxon dataset. We show average (over 20 replicates) quartet scores (sum of the weights of the satisfied quartets) of various methods, total weight of the quartets in the input gene trees, and their respective ratios.

| Model Condition | Tree | Quartet score | Total weight (input quartets) | Proportion (%) |
| --- | --- | --- | --- | --- |
| 100gene-100bp | Model Tree | 69307.4 | 136500.0 | 50.775 |
|  | wQFM | 69932.3 | 136500.0 | 51.232 |
|  | wQMC | 69930.0 | 136500.0 | 51.231 |
|  | ASTRAL | 69933.8 | 136500.0 | 51.234 |
| 100gene-1000bp | Model Tree | 82099.2 | 136500.0 | 60.146 |
|  | wQFM | 82166.6 | 136500.0 | 60.195 |
|  | wQMC | 82166.1 | 136500.0 | 60.195 |
|  | ASTRAL | 82166.6 | 136500.0 | 60.195 |
| 1000gene-100bp | Model Tree | 690268.0 | 1365000.0 | 50.569 |
|  | wQFM | 693656.1 | 1365000.0 | 50.817 |
|  | wQMC | 693656.1 | 1365000.0 | 50.817 |
|  | ASTRAL | 693656.1 | 1365000.0 | 50.817 |
| 1000gene-1000bp | Model Tree | 817937.3 | 1365000.0 | 59.922 |
|  | wQFM | 818022.2 | 1365000.0 | 59.928 |
|  | wQMC | 818022.2 | 1365000.0 | 59.928 |
|  | ASTRAL | 818022.2 | 1365000.0 | 59.928 |
| 100gene-true | Model Tree | 84634.9 | 136500.0 | 62.004 |
|  | wQFM | 84634.9 | 136500.0 | 62.004 |
|  | wQMC | 84634.9 | 136500.0 | 62.004 |
|  | ASTRAL | 84634.9 | 136500.0 | 62.004 |
| 1000gene-true | Model Tree | 844184.3 | 1365000.0 | 61.845 |
|  | wQFM | 844184.3 | 1365000.0 | 61.845 |
|  | wQMC | 844184.3 | 1365000.0 | 61.845 |
|  | ASTRAL | 844184.3 | 1365000.0 | 61.845 |

**Table S2:** Quartet scores on 11-taxon dataset. We show average (over 20 replicates) quartet scores (sum of the weights of the satisfied quartets) of the true and estimated trees, total weight of the quartets in the input gene trees, and their respective ratios.

| Model Condition | Tree | Quartet score | Total weight (input quartets) | Proportion (%) |
| --- | --- | --- | --- | --- |
| true-5genes | Model Tree | 1528.0 | 1650.0 | 92.606 |
|  | wQFM | 1536.8 | 1650.0 | 93.139 |
|  | wQMC | 1536.8 | 1650.0 | 93.139 |
|  | ASTRAL | 1536.8 | 1650.0 | 93.139 |
| true-15genes | Model Tree | 4619.5 | 4950.0 | 93.323 |
|  | wQFM | 4637.9 | 4950.0 | 93.695 |
|  | wQMC | 4637.9 | 4950.0 | 93.695 |
|  | ASTRAL | 4637.9 | 4950.0 | 93.695 |
| true-25genes | Model Tree | 7708.25 | 8250.0 | 93.433 |
|  | wQFM | 7708.25 | 8250.0 | 93.433 |
|  | wQMC | 7708.45 | 8250.0 | 93.436 |
|  | ASTRAL | 7708.45 | 8250.0 | 93.436 |
| true-50genes | Model Tree | 15412.55 | 16500.0 | 93.409 |
|  | wQFM | 15412.55 | 16500.0 | 93.409 |
|  | wQMC | 15412.55 | 16500.0 | 93.409 |
|  | ASTRAL | 15412.55 | 16500.0 | 93.409 |
| true-100genes | Model Tree | 30844.25 | 33000.0 | 93.467 |
|  | wQFM | 30844.25 | 33000.0 | 93.467 |
|  | wQMC | 30844.25 | 33000.0 | 93.467 |
|  | ASTRAL | 30844.25 | 33000.0 | 93.467 |
| estimated-5genes | Model Tree | 1321.3 | 1650.0 | 80.079 |
|  | wQFM | 1355.5 | 1650.0 | 82.152 |
|  | wQMC | 1355.05 | 1650.0 | 82.124 |
|  | ASTRAL | 1357.4 | 1650.0 | 82.267 |
| estimated-15genes | Model Tree | 4042.9 | 4950.0 | 81.675 |
|  | wQFM | 4068.25 | 4950.0 | 82.187 |
|  | wQMC | 4068.25 | 4950.0 | 82.187 |
|  | ASTRAL | 4068.25 | 4950.0 | 82.187 |
| estimated-25genes | Model Tree | 6679.45 | 8250.0 | 80.963 |
|  | wQFM | 6695.4 | 8250.0 | 81.156 |
|  | wQMC | 6695.4 | 8250.0 | 81.156 |
|  | ASTRAL | 6695.4 | 8250.0 | 81.156 |
| estimated-50genes | Model Tree | 13362.85 | 16500.0 | 80.987 |
|  | wQFM | 13383.1 | 16500.0 | 81.11 |
|  | wQMC | 13383.1 | 16500.0 | 81.11 |
|  | ASTRAL | 13383.1 | 16500.0 | 81.11 |
| estimated-100genes | Model Tree | 26686.55 | 33000.0 | 80.868 |
|  | wQFM | 26702.05 | 33000.0 | 80.915 |
|  | wQMC | 26702.05 | 33000.0 | 80.915 |
|  | ASTRAL | 26705.4 | 33000.0 | 80.925 |

**Table S3:** Quartet scores on 37-taxon mammalian simulated dataset. We show average (over 20 replicates) quartet scores (sum of the weights of the satisfied quartets) of the true and estimated trees, total weight of the quartets in the input gene trees, and their respective ratios. Various model conditions are defined by different ILS levels (1X, 0.5X, 2X), numbers of genes (100g, 200g, etc.) and sequence legths (500b, 100b, etc.).

| Model Condition | Tree | Quartet score | Total weight (input quartets) | Proportion (%) |
| --- | --- | --- | --- | --- |
| 1X-100g-500b | Model Tree | 5630260.55 | 6604500.0 | 85.249 |
|  | wQFM | 5635618.65 | 6604500.0 | 85.33 |
|  | wQMC | 5636032.85 | 6604500.0 | 85.336 |
|  | ASTRAL | 5636032.85 | 6604500.0 | 85.336 |
| 1X-200g-1000b | Model Tree | 11584969.95 | 13209000.0 | 87.705 |
|  | wQFM | 11586529.55 | 13209000.0 | 87.717 |
|  | wQMC | 11586641.45 | 13209000.0 | 87.718 |
|  | ASTRAL | 11586641.45 | 13209000.0 | 87.718 |
| 1X-200g-1500b | Model Tree | 11657249.9 | 13209000.0 | 88.252 |
|  | wQFM | 11658309.05 | 13209000.0 | 88.26 |
|  | wQMC | 11658432.55 | 13209000.0 | 88.261 |
|  | ASTRAL | 11658445.25 | 13209000.0 | 88.261 |
| 1X-200g-250b | Model Tree | 10557321.6 | 13209000.0 | 79.925 |
|  | wQFM | 10560880.15 | 13209000.0 | 79.952 |
|  | wQMC | 10561734.4 | 13209000.0 | 79.959 |
|  | ASTRAL | 10561825.45 | 13209000.0 | 79.959 |
| 1X-200g-true | Model Tree | 11744078.75 | 13209000.0 | 88.91 |
|  | wQFM | 11746143.2 | 13209000.0 | 88.925 |
|  | wQMC | 11746177.35 | 13209000.0 | 88.926 |
|  | ASTRAL | 11746203.5 | 13209000.0 | 88.926 |
| 1X-25g-500b | Model Tree | 1410250.7 | 1651125.0 | 85.412 |
|  | wQFM | 1414463.05 | 1651125.0 | 85.667 |
|  | wQMC | 1414450.05 | 1651125.0 | 85.666 |
|  | ASTRAL | 1414559.25 | 1651125.0 | 85.672 |
| 1X-400g-500b | Model Tree | 22531906.2 | 26418000.0 | 85.29 |
|  | wQFM | 22533922.6 | 26418000.0 | 85.298 |
|  | wQMC | 22534424.55 | 26418000.0 | 85.3 |
|  | ASTRAL | 22534424.55 | 26418000.0 | 85.3 |
| 1X-50g-500b | Model Tree | 2816708.35 | 3302250.0 | 85.297 |
|  | wQFM | 2820861.25 | 3302250.0 | 85.422 |
|  | wQMC | 2820892.25 | 3302250.0 | 85.423 |
|  | ASTRAL | 2821123.85 | 3302250.0 | 85.43 |
| 1X-800g-500b | Model Tree | 45096639.8 | 52836000.0 | 85.352 |
|  | wQFM | 45096766.4 | 52836000.0 | 85.352 |
|  | wQMC | 45096874.45 | 52836000.0 | 85.353 |
|  | ASTRAL | 45096874.45 | 52836000.0 | 85.353 |
| 0.5X-200g-500b | Model Tree | 10003929.55 | 13209000.0 | 75.736 |
|  | wQFM | 10006290.4 | 13209000.0 | 75.754 |
|  | wQMC | 10006510.55 | 13209000.0 | 75.755 |
|  | ASTRAL | 10007425.55 | 13209000.0 | 75.762 |
| 2X-200g-500b | Model Tree | 11943759.05 | 13205697.75 | 90.444 |
|  | wQFM | 11944321.3 | 13205697.75 | 90.448 |
|  | wQMC | 11944654.4 | 13205697.75 | 90.451 |
|  | ASTRAL | 11944654.4 | 13205697.75 | 90.451 |
| 1X-200g-500b | Model Tree | 11266716.15 | 13209000.0 | 85.296 |
|  | wQFM | 11270452.0 | 13209000.0 | 85.324 |
|  | wQMC | 11270657.55 | 13209000.0 | 85.326 |
|  | ASTRAL | 11270657.55 | 13209000.0 | 85.326 |

**Table S4:** Quartet scores on 48-taxon simulated avian dataset. We show average (over 20 replicates) quartet scores (sum of the weights of the satisfied quartets) of the true and estimated trees, total weight of the quartets in the input gene trees, and their respective ratios. Various model conditions are defined by different ILS levels (1X, 0.5X, 2X), numbers of genes (100g, 200g, etc.) and sequence legths (500b, 100b, etc.).

| Model Condition | Tree | Quartet score | Total weight (input quartets) | Proportion (%) |
| --- | --- | --- | --- | --- |
| 0.5X-1000g-500b | Model Tree | 110526512.9 | 194580000.0 | 56.803 |
|  | wQFM | 110133829.9 | 194580000.0 | 56.601 |
|  | wQMC | 109888348.3 | 194580000.0 | 56.475 |
|  | ASTRAL | 110583148.55 | 194580000.0 | 56.832 |
| 1X-1000g-500b | Model Tree | 122945091.0 | 194580000.0 | 63.185 |
|  | wQFM | 122897544.85 | 194580000.0 | 63.16 |
|  | wQMC | 122865927.3 | 194580000.0 | 63.144 |
|  | ASTRAL | 123024937.95 | 194580000.0 | 63.226 |
| 2X-1000g-500b | Model Tree | 131567523.05 | 194580000.0 | 67.616 |
|  | wQFM | 131455003.5 | 194580000.0 | 67.558 |
|  | wQMC | 131457328.1 | 194580000.0 | 67.56 |
|  | ASTRAL | 131631393.9 | 194580000.0 | 67.649 |
| 1X-50g-500b | Model Tree | 6109255.45 | 9729000.0 | 62.794 |
|  | wQFM | 6203890.95 | 9729000.0 | 63.767 |
|  | wQMC | 6209354.0 | 9729000.0 | 63.823 |
|  | ASTRAL | 6204883.7 | 9729000.0 | 63.777 |
| 1X-100g-500b | Model Tree | 12250259.15 | 19458000.0 | 62.957 |
|  | wQFM | 12343389.35 | 19458000.0 | 63.436 |
|  | wQMC | 12342740.1 | 19458000.0 | 63.433 |
|  | ASTRAL | 12349310.6 | 19458000.0 | 63.466 |
| 1X-200g-500b | Model Tree | 24553380.5 | 38916000.0 | 63.093 |
|  | wQFM | 24618478.25 | 38916000.0 | 63.261 |
|  | wQMC | 24618794.95 | 38916000.0 | 63.261 |
|  | ASTRAL | 24632823.8 | 38916000.0 | 63.297 |
| 1X-500g-500b | Model Tree | 61438944.15 | 97290000.0 | 63.15 |
|  | wQFM | 61463222.85 | 97290000.0 | 63.175 |
|  | wQMC | 61440254.85 | 97290000.0 | 63.152 |
|  | ASTRAL | 61528828.05 | 97290000.0 | 63.243 |

#### 4.5 Wilcoxon Signed-Rank Test

Tables S5 and S6 show the *p*-values using the Wilcoxon signed-rank test (with *α* = 0.05) on various model conditions of 11-, 15-, 37-, and 48-taxon datasets, comparing wQFM with wQMC and ASTRAL, respectively.

**Table S5:** Statistical significance of the differences between **wQFM** and **wQMC** in 11-, 15-, 37- and 48-taxon datasets. The p-values indicating statistically significant differences (i.e., *p* ≤ 0.05) are shown in bold.

| *Dataset* | *Model condition* |  |  | *p-value* |
| --- | --- | --- | --- | --- |
|  | *No. of genes* | *Sequence len.* | *ILS Level* |  |
| **11-taxon** | 5 g | true | high | **0.0255** |
|  | 15 g | true | high | 0.1572 |
|  | 25 g | true | high | 0.3173 |
|  | 5 g | estimated | high | 0.3049 |
| **15-taxon** | 100 g | 100 bp | high | 0.3173 |
|  | 100 g | 1000 bp | high | 0.3173 |
| **37-taxon** | 25 g | 500 bp | 1X | 0.9814 |
|  | 50 g | 500 bp | 1X | 0.6744 |
|  | 100 g | 500 bp | 1X | 0.9444 |
|  | 200 g | 500 bp | 1X | **0.0455** |
|  | 200 g | 500 bp | 0.5X | 0.1250 |
|  | 200 g | 500 bp | 2X | 0.3173 |
|  | 400 g | 500 bp | 1X | 0.3173 |
|  | 800 g | 500 bp | 1X | 0.1572 |
|  | 200 g | 250 bp | 1X | **0.0033** |
|  | 200 g | 1000 bp | 1X | 0.0832 |
|  | 200 g | 1500 bp | 1X | **0.0458** |
|  | 200 g | true | 1X | 0.0832 |
| **48-taxon** | 1000 g | 500 bp | 0.5X | **0.0011** |
|  | 1000 g | 500 bp | 1X | **0.0381** |
|  | 1000 g | 500 bp | 2X | 0.8316 |
|  | 50g | 500 bp | 1X | **0.0235** |
|  | 100g | 500 bp | 1X | **0.0064** |
|  | 200g | 500 bp | 1X | **0.0002** |
|  | 500g | 500 bp | 1X | **0.0249** |

**Table S6:** Statistical significance of the differences between **wQFM** and **ASTRAL** in 11-, 15-, 37- and 48-taxon datasets. The p-values indicating statistically significant differences (i.e., *p* ≤ 0.05) are shown in bold.

| *Dataset* | *Model condition* |  |  | *p-value* |
| --- | --- | --- | --- | --- |
|  | *No. of genes* | *Sequence len.* | *ILS Level* |  |
| **11-taxon** | 5 g | true | high | 0.0832 |
|  | 15 g | true | high | 0.1572 |
|  | 25 g | true | high | 0.3173 |
|  | 5 g | estimated | high | 0.1797 |
|  | 100 g | estimated | high | 0.3173 |
| **15-taxon** | 100 g | 100 bp | high | 0.3173 |
| **37-taxon** | 25 g | 500 bp | 1X | 0.9354 |
|  | 50 g | 500 bp | 1X | 0.3901 |
|  | 100 g | 500 bp | 1X | 0.9444 |
|  | 200 g | 500 bp | 1X | **0.0455** |
|  | 200 g | 500 bp | 0.5X | 0.3750 |
|  | 200 g | 500 bp | 2X | 0.3173 |
|  | 400 g | 500 bp | 1X | 0.3173 |
|  | 800 g | 500 bp | 1X | 0.1572 |
|  | 200 g | 250 bp | 1X | **0.0157** |
|  | 200 g | 1000 bp | 1X | 0.0832 |
|  | 200 g | 1500 bp | 1X | 0.0832 |
|  | 200 g | true | 1X | 0.1572 |
| **48-taxon** | 1000 g | 500 bp | 0.5X | **0.0016** |
|  | 1000 g | 500 bp | 1X | **0.0003** |
|  | 1000 g | 500 bp | 2X | **0.0001** |
|  | 50 g | 500 bp | 1X | 0.2435 |
|  | 100 g | 500 bp | 1X | 0.9399 |
|  | 200 g | 500 bp | 1X | 0.5796 |
|  | 500 g | 500 bp | 1X | **0.0102** |

## References

[1] J F C Kingman. The coalescent. Stoch Proc Appl, 13:235–248, 1982.

[2] W. P. Maddison. Gene trees in species trees. Systematic Biology, 46:523–536, 1997.

[3] J H Degnan and N A Rosenberg. Gene tree discordance, phylogenetic inference and the multispecies coalescent. Trends Ecology Evolution, 26(6), 2009.

[4] S V Edwards. Is a new and general theory of molecular systematics emerging? Evolution, 63(1):1–19, 2009.

[5] Sebastien Roch and Mike Steel. Likelihood-based tree reconstruction on a concatenation of aligned sequence data sets can be statistically inconsistent. Theoretical population biology, 100:56–62, 2015.

[6] L S Kubatko and J H Degnan. Inconsistency of phylogenetic estimates from concatenated data under coalescence. Syst Biol, 56:17, 2007.

[7] Scott V Edwards, Liang Liu, and Dennis K Pearl. High-resolution species trees without concatenation. Proceedings of the National Academy of Sciences, 104(14):5936–5941, 2007.

[8] A D Leaché and B Rannala. The accuracy of species tree estimation under simulation: a comparison of methods. Syst Biol, 60(2):126–137, 2011.

[9] Michael DeGiorgio and James H Degnan. Fast and consistent estimation of species trees using supermatrix rooted triples. Molecular biology and evolution, 27(3):552–569, 2009.

[10] James H Degnan. Anomalous unrooted gene trees. Systematic biology, 62(4):574–590, 2013.

[11] Md Shamsuzzoha Bayzid and Tandy Warnow. Naive binning improves phylogenomic analyses. Bioinformatics, 29(18):2277–2284, 2013.

[12] J Heled and A J Drummond. Bayesian inference of species trees from multilocus data. Mol Biol Evol, 27:570–580, 2010.

[13] E Mossel and S Roch. Incomplete lineage sorting: consistent phylogeny estimation from multiple loci. IEEE Comp Biol Bioinform, 7(1):166–171, 2011.

[14] L S Kubatko, B C Carstens, and L L Knowles. Stem: Species tree estimation using maximum likelihood for gene trees under coalescence. Bioinf, 25:971–973, 2009.

[15] Siavash Mirarab, Rezwana Reaz, Md S Bayzid, Théo Zimmermann, M Shel Swenson, and Tandy Warnow. ASTRAL: genome-scale coalescent-based species tree estimation. Bioinformatics, 30(17):i541–i548, 2014.

[16] Liang Liu, Lili Yu, and Scott V Edwards. A maximum pseudo-likelihood approach for estimating species trees under the coalescent model. BMC Evolutinary Biology, 10:302, 2010.

[17] Liang Liu and Lili Yu. Estimating species trees from unrooted gene trees. Systematic Biology, 60(5):661–667, 2011.

[18] B Larget, S K Kotha, C N Dewey, and C Ané. BUCKy: Gene tree/species tree reconciliation with the Bayesian concordance analysis. Bioinf, 26(22):2910–2911, 2010.

[19] David Bryant, Remco Bouckaert, Joseph Felsenstein, Noah A Rosenberg, and Arindam RoyChoudhury. Inferring species trees directly from biallelic genetic markers: bypassing gene trees in a full coalescent analysis. Molecular biology and evolution, 29(8):1917–1932, 2012.

[20] Julia Chifman and Laura Kubatko. Quartet from snp data under the coalescent model. Bioinformatics, 30(23):3317–3324, 2014.

[21] Liang Liu, Lili Yu, Dennis K Pearl, and Scott V Edwards. Estimating species phylogenies using coalescence times among sequences. Systematic biology, 58(5):468–477, 2009.

[22] Pranjal Vachaspati and Tandy Warnow. Astrid: accurate species trees from internode distances. BMC genomics, 16(10):S3, 2015.

[23] Mazharul Islam, Kowshika Sarker, Trisha Das, Rezwana Reaz, and Md Shamsuzzoha Bayzid. Stelar: A statistically consistent coalescent-based species tree estimation method by maximizing triplet consistency. BMC genomics, 21(1):1–13, 2020.

[24] Rezwana Reaz, Md Shamsuzzoha Bayzid, and M Sohel Rahman. Accurate phylogenetic tree reconstruction from quartets: A heuristic approach. PLoS One, 9(8):e104008, 2014.

[25] K Strimmer and A von Haeseler. Quartet puzzling: A quartet maximim-likelihood method for reconstructing tree topologies. Molecular Biology and Evolution, 13(7):964–969, 1996.

[26] Heiko A Schmidt, Korbinian Strimmer, Martin Vingron, and Arndt von Haeseler. Tree-puzzle: maximum likelihood phylogenetic analysis using quartets and parallel computing. Bioinformatics, 18(3):502–504, 2002.

[27] V Ranwez and O Gascuel. Quartet-Based phylogenetic inference: Improvements and limits. Mol Biol Evol, 18(6):1103–1116, June 2001.

[28] Bin Ma, Lei Xin, and Kaizhong Zhang. A new quartet approach for reconstructing phylogenetic trees: quartet joining method. Journal of combinatorial optimization, 16(3):293–306, 2008.

[29] S. Snir and S. Rao. Quartets MaxCut: a divide and conquer quartets algorithm. IEEE/ACM Trans. Comput. Biol. Bioinform., 7(4):704–718, 2010.

[30] Eliran Avni, Reuven Cohen, and Sagi Snir. Weighted quartets phylogenetics. Systematic biology, 64(2):233–242, 2015.

[31] B. Chor and T. Tuller. Maximum likelihood of evolutionary trees: hardness and approximation. Bioinformatics, 21(1):97–106, 2005.

[32] Barbara R Holland, Peter D Jarvis, and Jeremy G Sumner. Low-parameter phylogenetic inference under the general markov model. Systematic biology, 62(1):78–92, 2013.

[33] David Bryant and Mike Steel. Constructing optimal trees from quartets. Journal of Algorithms, 38(1):237–259, 2001.

[34] Huateng Huang, Qixin He, Laura S Kubatko, and L Lacey Knowles. Sources of error inherent in species-tree estimation: impact of mutational and coalescent effects on accuracy and implications for choosing among different methods. Systematic Biology, 59(5):573–583, 2010.

[35] John Gatesy and Mark S Springer. Phylogenetic analysis at deep timescales: unreliable gene trees, bypassed hidden support, and the coalescence/concatalescence conundrum. Molecular phylogenetics and evolution, 80:231–266, 2014.

[36] M.A. Steel. The complexity of reconstructing trees from qualitative characters and subtrees. Journal of Classification, 9:91–116, 1992.

[37] Vincent Berry and Olivier Gascuel. Inferring evolutionary trees with strong combinatorial evidence. Theoretical Computer Science, 240:271–298, 2001.

[38] Victor C Mason, Gang Li, Patrick Minx, Jürgen Schmitz, Gennady Churakov, Liliya Doronina, Amanda D Melin, Nathaniel J Dominy, Norman TL Lim, Mark S Springer, et al. Genomic analysis reveals hidden biodiversity within colugos, the sister group to primates. Science advances, 2(8):e1600633, 2016.

[39] Hernán Vázquez-Miranda, Josie A Griffin, Jay M Sheppard, Jordan M Herman, Octavio Rojas-Soto, and Robert M Zink. Morphological and molecular evolution and their consequences for conservation and taxonomy in the le conte’s thrasher toxostoma lecontei. Journal of Avian Biology, 48(7):941–954, 2017.

[40] Anne D Yoder, C Ryan Campbell, Marina B Blanco, Mario Dos Reis, Jörg U Ganzhorn, Steven M Goodman, Kelsie E Hunnicutt, Peter A Larsen, Peter M Kappeler, Rodin M Rasoloarison, et al. Geogenetic patterns in mouse lemurs (genus microcebus) reveal the ghosts of madagascar’s forests past. Proceedings of the National Academy of Sciences, 113(29):8049–8056, 2016.

[41] Richard GJ Hodel, L Lacey Knowles, Stuart F McDaniel, Adam C Payton, Jordan F Dunaway, Pamela S Soltis, and Douglas E Soltis. Terrestrial species adapted to sea dispersal: Differences in propagule dispersal of two caribbean mangroves. Molecular ecology, 27(22):4612–4626, 2018.

[42] Kato Dai-ichiro, Suzuki Hirobumi, Tsuruta Atsuhiro, Maeda Juri, Hayashi Yoshinobu, Arima Kazunari, Ito Yuji, and Nagano Yukio. Evaluation of the population structure and phylogeography of the japanese genji firefly, luciola cruciata, at the nuclear dna level using rad-seq analysis. Scientific Reports (Nature Publisher Group), 10(1), 2020.

[43] Peter A Hosner, Edward L Braun, and Rebecca T Kimball. Rapid and recent diversification of curassows, guans, and chachalacas (galliformes: Cracidae) out of mesoamerica: Phylogeny inferred from mitochondrial, intron, and ultraconserved element sequences. Molecular phylogenetics and evolution, 102:320–330, 2016.

[44] Thomas J Devitt, April M Wright, David C Cannatella, and David M Hillis. Species delimitation in endangered groundwater salamanders: Implications for aquifer management and biodiversity conservation. Proceedings of the National Academy of Sciences, 116(7):2624–2633, 2019.

[45] Milan Malinsky, Hannes Svardal, Alexandra M Tyers, Eric A Miska, Martin J Genner, George F Turner, and Richard Durbin. Whole-genome sequences of malawi cichlids reveal multiple radiations interconnected by gene flow. Nature ecology & evolution, 2(12):1940–1955, 2018.

[46] Nazifa Ahmed Moumi, Badhan Das, Zarin Tasnim Promi, Nishat Anjum Bristy, and Md Shamsuzzoha Bayzid. Quartet-based inference of cell differentiation trees from chip-seq histone modification data. PloS one, 14(9), 2019.

[47] Dai-ichiro Kato, Hirobumi Suzuki, Atsuhiro Tsuruta, Juri Maeda, Yoshinobu Hayashi, Kazunari Arima, Yuji Ito, and Yukio Nagano. Evaluation of the population structure and phylogeography of the japanese genji firefly, luciola cruciata, at the nuclear dna level using rad-seq analysis. Scientific reports, 10(1):1–12, 2020.

[48] Julia Chifman and Laura Kubatko. Identifiability of the unrooted species tree topology under the coalescent model with time-reversible substitution processes, site-specific rate variation, and invariable sites. Journal of theoretical biology, 374:35–47, 2015.

[49] D.L. Swofford. PAUP*: Phylogenetic analysis using parsimony (* and other methods). Ver. 4. Sinauer Associates, Sunderland, Massachusetts, 2002.

[50] Jed Chou, Ashu Gupta, Shashank Yaduvanshi, Ruth Davidson, Mike Nute, Siavash Mirarab, and Tandy Warnow. A comparative study of svdquartets and other coalescent-based species tree estimation methods. BMC genomics, 16(10):S2, 2015.

[51] Erich D Jarvis, Siavash Mirarab, Andre J Aberer, Bo Li, Peter Houde, Cai Li, Simon YW Ho, Brant C Faircloth, Benoit Nabholz, Jason T Howard, et al. Wholegenome analyses resolve early branches in the tree of life of modern birds. Science, 346(6215):1320–1331, 2014.

[52] C Anée, B Larget, D A Baum, S D Smith, and A Rokas. Bayesian estimation of concordance among gene trees. Mol Biol Evol, 24:412–426, 2007.

[53] P Erdos, M Steel, L Szekely, and T Warnow. A few logs suffice to build (almost) all trees (i). Random Structures and Algorithms, 14:153–184, 1999.

[54] T. Warnow J. Yang. Fast and accurate methods for phylogenomic analyses. volume 12(Suppl 9), 2011.

[55] T Jiang, P Kearney, and M Li. A polynomial-time approximation scheme for inferring evolutionary trees from quartet topologies and its applications. SIAM J. Comput.,30(6):1924–1961, 2001.

[56] Siavash Mirarab, Md Shamsuzzoha Bayzid, Bastien Boussau, and Tandy Warnow. Statistical binning enables an accurate coalescent-based estimation of the avian tree. Science, 346(6215):1250463, 2014.

[57] Y Chung and C Ané. Comparing two Bayesian methods for gene tree/species tree reconstruction: A simulation with incomplete lineage sorting and horizontal gene transfer. Syst Biol, 60(3):261–275, 2011.

[58] Siavash Mirarab and Tandy Warnow. Astral-ii: coalescent-based species tree estimation with many hundreds of taxa and thousands of genes. Bioinformatics, 31(12):i44–i52, 2015.

[59] Sen Song, Liang Liu, Scott V Edwards, and Shaoyuan Wu. Resolving conflict in eutherian mammal phylogeny using phylogenomics and the multispecies coalescent model. Proceedings of the National Academy of Sciences, 109(37):14942–14947, 2012.

[60] Ylenia Chiari, Vincent Cahais, Nicolas Galtier, and Frédéric Delsuc. Phylogenomic analyses support the position of turtles as the sister group of birds and crocodiles (archosauria). Bmc Biology, 10(1):65, 2012.

[61] Zhenxiang Xi, Liang Liu, Joshua S Rest, and Charles C Davis. Coalescent versus concatenation methods and the placement of amborella as sister to water lilies. Systematic biology, 63(6):919–932, 2014.

[62] Chao Zhang, Maryam Rabiee, Erfan Sayyari, and Siavash Mirarab. Astral-iii: polynomial time species tree reconstruction from partially resolved gene trees. BMC bioinformatics, 19(6):153, 2018.

[63] Ruth Davidson, Pranjal Vachaspati, Siavash Mirarab, and Tandy Warnow. Phylogenomic species tree estimation in the presence of incomplete lineage sorting and horizontal gene transfer. BMC genomics, 16(S10):S1, 2015.

[64] D.F. Robinson and L.R. Foulds. Comparison of phylogenetic trees. Math. Biosci., 53:131–147, 1981.

[65] Ishrat Tanzila Farah, Md Muktadirul Islam, Kazi Tasnim Zinat, Atif Hasan Rahman, and Md Shamsuzzoha Bayzid. Phylogenomic terraces: presence and implication in species tree estimation from gene trees. bioRxiv, 2020.

[66] Alexey M Kozlov, Andre J Aberer, and Alexandros Stamatakis. Examl version 3: a tool for phylogenomic analyses on supercomputers. Bioinformatics, 31(15):2577–2579, 2015.

[67] Md Shamsuzzoha Bayzid, Siavash Mirarab, Bastien Boussau, and Tandy Warnow. Weighted statistical binning: enabling statistically consistent genome-scale phylogenetic analyses. PLoS One, 10(6), 2015.

[68] Per GP Ericson, Cajsa L Anderson, Tom Britton, Andrzej Elzanowski, Ulf S Johansson, Mari Källersjö, Jan I Ohlson, Thomas J Parsons, Dario Zuccon, and Gerald Mayr. Diversification of neoaves: integration of molecular sequence data and fossils. Biology letters, 2(4):543–547, 2006.

[69] Shannon J Hackett, Rebecca T Kimball, Sushma Reddy, Rauri CK Bowie, Edward L Braun, Michael J Braun, Jena L Chojnowski, W Andrew Cox, Kin-Lan Han, John Harshman, et al. A phylogenomic study of birds reveals their evolutionary history. science, 320(5884):1763–1768, 2008.

[70] Rebecca T Kimball, Ning Wang, Victoria Heimer-McGinn, Carly Ferguson, and Edward L Braun. Identifying localized biases in large datasets: a case study using the avian tree of life. Molecular Phylogenetics and Evolution, 69(3):1021–1032, 2013.

[71] Joel Cracraft. Avian higher-level relationships and classification: nonpasseriforms. The Howard and Moore complete checklist of the birds of the world, 1:21–47, 2013.

[72] Siavash Mirarab, Md Shamsuzzoha Bayzid, and Tandy Warnow. Evaluating summary methods for multilocus species tree estimation in the presence of incomplete lineage sorting. Systematic Biology, 65(3):366–380, 2014.

[73] Jan E Janeĉka, Webb Miller, Thomas H Pringle, Frank Wiens, Annette Zitzmann, Kristofer M Helgen, Mark S Springer, and William J Murphy. Molecular and genomic data identify the closest living relative of primates. Science, 318(5851):792–794, 2007.

[74] Vikas Kumar, Björn M Hallström, and Axel Janke. Coalescent-based genome analyses resolve the early branches of the euarchontoglires. PLoS One, 8(4):e60019, 2013.

[75] B. Boussau, G. J. Szöllősi, L. Duret, M. Gouy, E. Tannier, and V. Daubin. Genomescale coestimation of species and gene trees. Genome Research, 23(2):323–330, 2013.

[76] HU Jingyang, ZHANG Yaping, and YU Li. Summary of laurasiatheria (mammalia) phylogeny. Zoological Research, 33(E5-6):65–74, 2012.

[77] Andrew F Hugall, Ralph Foster, and Michael SY Lee. Calibration choice, rate smoothing, and the pattern of tetrapod diversification according to the long nuclear gene rag-1. Systematic biology, 56(4):543–563, 2007.

[78] Naoyuki Iwabe, Yuichiro Hara, Yoshinori Kumazawa, Kaori Shibamoto, Yumi Saito, Takashi Miyata, and Kazutaka Katoh. Sister group relationship of turtles to the bird-crocodilian clade revealed by nuclear dna–coded proteins. Molecular Biology and Evolution, 22(4):810–813, 2004.

[79] Norman J Wickett, Siavash Mirarab, Nam Nguyen, Tandy Warnow, Eric Carpenter, Naim Matasci, Saravanaraj Ayyampalayam, Michael S Barker, J Gordon Burleigh, Matthew A Gitzendanner, et al. Phylotranscriptomic analysis of the origin and early diversification of land plants. Proceedings of the National Academy of Sciences, 111(45):E4859–E4868, 2014.

[80] Ning Zhang, Liping Zeng, Hongyan Shan, and Hong Ma. Highly conserved low-copy nuclear genes as effective markers for phylogenetic analyses in angiosperms. New Phytologist, 195(4):923–937, 2012.

[81] Bryan T Drew, Brad R Ruhfel, Stephen A Smith, Michael J Moore, Barbara G Briggs, Matthew A Gitzendanner, Pamela S Soltis, and Douglas E Soltis. Another look at the root of the angiosperms reveals a familiar tale. Systematic Biology, 63(3):368–382, 2014.

[82] Vadim V Goremykin, Svetlana V Nikiforova, Patrick J Biggs, Bojian Zhong, Peter Delange, William Martin, Stefan Woetzel, Robin A Atherton, Patricia A Mcle-nachan, and Peter J Lockhart. The evolutionary root of flowering plants. Systematic Biology, 62(1):50–61, 2013.

## References

[1] Julia Chifman and Laura Kubatko. Quartet from snp data under the coalescent model. Bioinformatics, 30(23):3317–3324, 2014.

